# Climate warming reduces the speed and predictability of polygenic adaptation to salinity decline

**DOI:** 10.64898/2026.08.01.742181

**Authors:** Zhenyong Du, Alexander Taylor, Lydia Larsen, Samuel Lu, Carol Eunmi Lee

## Abstract

Climate change exposes populations to multiple stressors simultaneously, yet our understanding of how the addition of one stressor alters adaptation to another remains poor. As a result of climate change, high-latitude coastal habitats are experiencing rapid salinity decline, resulting in serious impacts on food webs and ocean circulation. Here, we examine how temperature increase impacts adaptation to salinity decline, in terms of the speed, genomic response, and repeatability of adaptation. We performed replicated Evolve-and-Resequence experiments over 20 to 25 generations in the model copepod *Eurytemora carolleeae* (Atlantic clade of the *E. affinis* species complex). Under salinity decline alone, replicate selection lines exhibited a polygenic response involving 66 selected haplotype blocks, with increasing parallelism among the replicate lines through Generation 20. Fitness (egg number) declined sharply over the first four generations but underwent full Evolutionary Rescue, recovering to ancestral levels by Generation 10. In contrast, imposing temperature increase on the salinity decline lines resulted in significantly lower parallelism among the selection lines, along with delayed and incomplete Evolutionary Rescue. Only 14% of selected SNPs were shared between the two selection regimes, and Gene Ontology analyses revealed largely distinct functional categories of genes under selection. These results show that adding warming to salinity decline can alter the genomic trajectory of salinity adaptation, slowing and impeding Evolutionary Rescue, and reducing the repeatability of polygenic responses. Our findings have direct implications for predicting evolutionary responses to realistic, multi-stressor climate change in high-latitude coastal ecosystems experiencing simultaneous ocean freshening and warming.

## Introduction

Populations facing rapid environmental change that exceeds their tolerance thresholds must adapt, migrate, or face extinction (Lee & Gelembiuk, 2008; Chevin et al., 2010; Tellier et al., 2024). Adaptation to environmental stressors often involves polygenic responses, with coordinated allele frequency shifts across hundreds or thousands of genomic loci (Höllinger et al., 2019; Barghi et al., 2020; Stern & Lee, 2020). A fundamental question in evolutionary biology is whether such polygenic responses are predictable, that is, whether independent populations exposed to the same selective pressure show parallel evolutionary responses (Gould, 1989; Blount et al., 2018; Bolnick et al., 2018). The degree of parallel evolution during replicate adaptive events provides direct insights into whether evolutionary pathways are constrained or labile (Conte et al., 2012; Stern, 2013), with profound implications for forecasting population persistence under climate change (Waldvogel et al., 2019; Bernatchez et al., 2024).

Recent Evolve-and-Resequence (E&R) studies have revealed a wide range of parallelism during polygenic adaptation, from remarkably repeatable responses to highly divergent and idiosyncratic evolutionary trajectories (Barghi et al., 2019; Otte et al., 2021a; Stern et al., 2022; Reid et al., 2023; Schlötterer, 2023; Forsberg et al., 2025; Du et al., 2026). We define “parallelism” here as the degree to which independent populations exposed to the same selective pressure evolve through allele frequency changes at the same genomic loci or linked haplotype blocks. For instance, in the copepod *Eurytemora affinis* species complex (Lee, 1999, 2000), an evolution experiment exhibited exceptionally high parallelism under salinity decline in the Europe clade of this complex (*E. affinis* proper, 15 chromosomes), with 50.8 to 79.5% of selected alleles shared across replicate lines (Stern et al., 2022; Du et al., 2026). Simulations of these data indicated that this high level of parallelism was consistent with positive synergistic epistasis driving coordinated allele frequency shifts across replicate selection lines. In a companion study, Du et al. (2026) showed that the Atlantic clade of the *E. affinis* complex (*E. carolleeae*, 4 chromosomes) exhibited substantially lower parallelism in response to salinity decline. This lower degree of parallelism was attributable to selection on large linked haplotype blocks inherited as units within a genome, likely arising from fewer chromosomes and lower recombination rates. These contrasting selection responses demonstrate that genome architecture, particularly chromosome number and recombination landscape, profoundly impacts the mechanism and predictability of polygenic adaptation to salinity decline (Lee, 2023, 2025; Du et al., 2026; Wirtz et al., 2026).

However, in nature, populations rarely face single environmental stressors in isolation. Climate change is driving simultaneous shifts in multiple abiotic variables, including temperature, salinity, pH, and oxygen concentration (Boyd et al., 2018; Lee et al., 2022; Röthig et al., 2023). High-latitude coastal waters are experiencing remarkably rapid transformations in these environmental variables. In particular, salinity is projected to decline by up to 5‰ (g/kg) in many high-latitude coastal habitats in the coming decades due to increased precipitation and enormous freshwater input from ice melt (Durack et al., 2012; Lavoie et al., 2015; Long & Perrie, 2015). The freshening of the North Atlantic is currently disrupting deep water formation and could eventually lead to the collapse of the Atlantic Meridional Overturning Circulation (AMOC), resulting in deep ocean anoxia and the demise of the Gulf Stream (Rahmstorf, 2024). At the same time, ocean warming could occur at rates of 2.5 to 4.7°C per century (Gröger et al., 2019). Despite the pervasiveness of such concurrent environmental shifts, most E&R studies of polygenic adaptation have not examined responses to the simultaneous effects of salinity and temperature (Hsu et al., 2024). In general, we understand little about how adding a second, co-occurring stressor affects the speed, genomic mechanisms, and repeatability of a well-characterized adaptive response, aside from a few case studies (Dam et al., 2021; Brennan et al., 2022). Thus, in this study, we examine how the rise in temperature impacts the well-studied population genomic response to salinity decline (Stern & Lee, 2020; Stern et al., 2022; Du et al., 2025) in the context of Evolve-and-Resequence (E&R) experiments.

Theoretical models predict that selection imposed by additional environmental stressors should reduce both the speed and predictability of adaptation through several mechanisms (Rose, 1982; Orr, 2000; Walsh & Blows, 2009; Tenaillon, 2014; Teplitsky et al., 2014; Castellano et al., 2016). For instance, Fisher’s geometric model demonstrates a “cost of complexity,” in which the rate of adaptation declines as the number of phenotypic dimensions under selection increases, because the probability that a random mutation simultaneously improves all traits diminishes (Orr, 2000; Tenaillon, 2014). In addition, multivariate genetic constraints can impede evolutionary responses in the adaptive direction even when individual traits harbor ample genetic variation (Walsh & Blows, 2009; Teplitsky et al., 2014). Selection on an additional stressor can also increase genetic redundancy by allowing a larger number of genetic combinations to contribute to adaptation. As the number of contributing loci grows, a greater proportion of them become functionally interchangeable, such that different replicate populations may recruit different subsets of loci, reducing parallelism among populations exposed to the same selection regime (Barghi et al., 2019; Barghi et al., 2020; Láruson et al., 2020). Additional effects, including the Hill–Robertson effect among simultaneously selected linked loci (Castellano et al., 2016) and antagonistic pleiotropy between traits adapted to different stressors (Rose, 1982), might further impede adaptation when more than one stressor is present.

Direct insights into genomic responses to the joint effects of salinity and temperature come from wild *E. affinis* proper populations spanning salinity and temperature gradients in the Baltic Sea and North Sea region. Across this range, *E. affinis* proper populations exhibited clear genomic signatures of selection in response to both environmental gradients, but the underlying genes involved were largely non-overlapping (Diaz et al., 2026). Correlations between single nucleotide polymorphisms (SNPs) associated with salinity versus temperature were low, with only 5 to 18% of significant SNPs shared between the two environmental variables. Ion transport-related functional categories dominated the salinity response, whereas the temperature response involved many SNPs but tended to show no enrichment of any particular functional category (Diaz et al., 2026).

These prior theoretical and empirical studies generate several specific predictions regarding how adding temperature increase should alter the salinity selection response. First of all, we predicted that adding warming to the salinity decline selection regime would impose a greater detrimental impact on fitness than salinity decline alone, thereby slowing population recovery and impeding “Evolutionary Rescue” (Gonzalez et al., 2013; Anciaux et al., 2018). Due to the differences in the physiological machinery involved in salinity versus temperature responses, adapting to both variables simultaneously would entail a higher fitness cost than adapting to salinity alone (deMayo et al., 2025). Second, we predicted that adding warming would broaden the genomic response and recruit a largely distinct set of loci, involving a greater number of small-effect candidate SNPs. The low correlation between salinity and temperature responses in the wild populations (see previous paragraph) suggests that temperature and salinity changes in an evolution experiment should each recruit largely distinct sets of genomic loci.

Finally, we predicted that adding temperature would reduce the parallelism among replicate selection lines, particularly the high levels of parallelism we observed previously for salinity decline alone (Stern & Lee, 2020; Stern et al., 2022; Du et al., 2026). As the temperature selection response appears to be more polygenic than the salinity response (Diaz et al., 2026), we would expect that temperature adaptation would involve a more redundant set of selected loci (Barghi et al., 2019, 2020).

This study therefore examined how temperature change, specifically warming, impacts the genomic response to salinity decline in replicate selection lines of the copepod *E. carolleeae* (Atlantic clade of the *E. affinis* species complex). Specifically, our goals were to determine how the addition of warming to the salinity decline selection regime impacted (1) the speed and completeness of fitness recovery (Evolutionary Rescue) associated with adaptation, (2) the genomic architecture of the selection response, including the number, trajectory, and chromosomal distribution of the loci under selection, (3) the extent of parallelism among replicate selection lines, and (4) the functional pathways of genes associated with adaptation.

To address these goals, we performed two complementary Evolve-and-Resequence (E&R) experiments using *E. carolleeae* (haploid chromosome number = 4; Fig. 1a). In the first selection regime, selection lines experienced salinity decline alone (S) from saline water (15‰) to fresh water (0‰) over 20 generations. In the second regime, selection lines experienced the same salinity decline (15‰ to 0‰) combined with temperature increase (ST; 12°C to 22°C, then maintained at 20°C) over 25 generations (Fig. 1b). Both experiments used the same source population from the St. Lawrence estuary at Baie de L’Isle Verte, Quebec, Canada, so that the two selection regimes differed only in the addition of temperature increase.

**Fig. 1.**
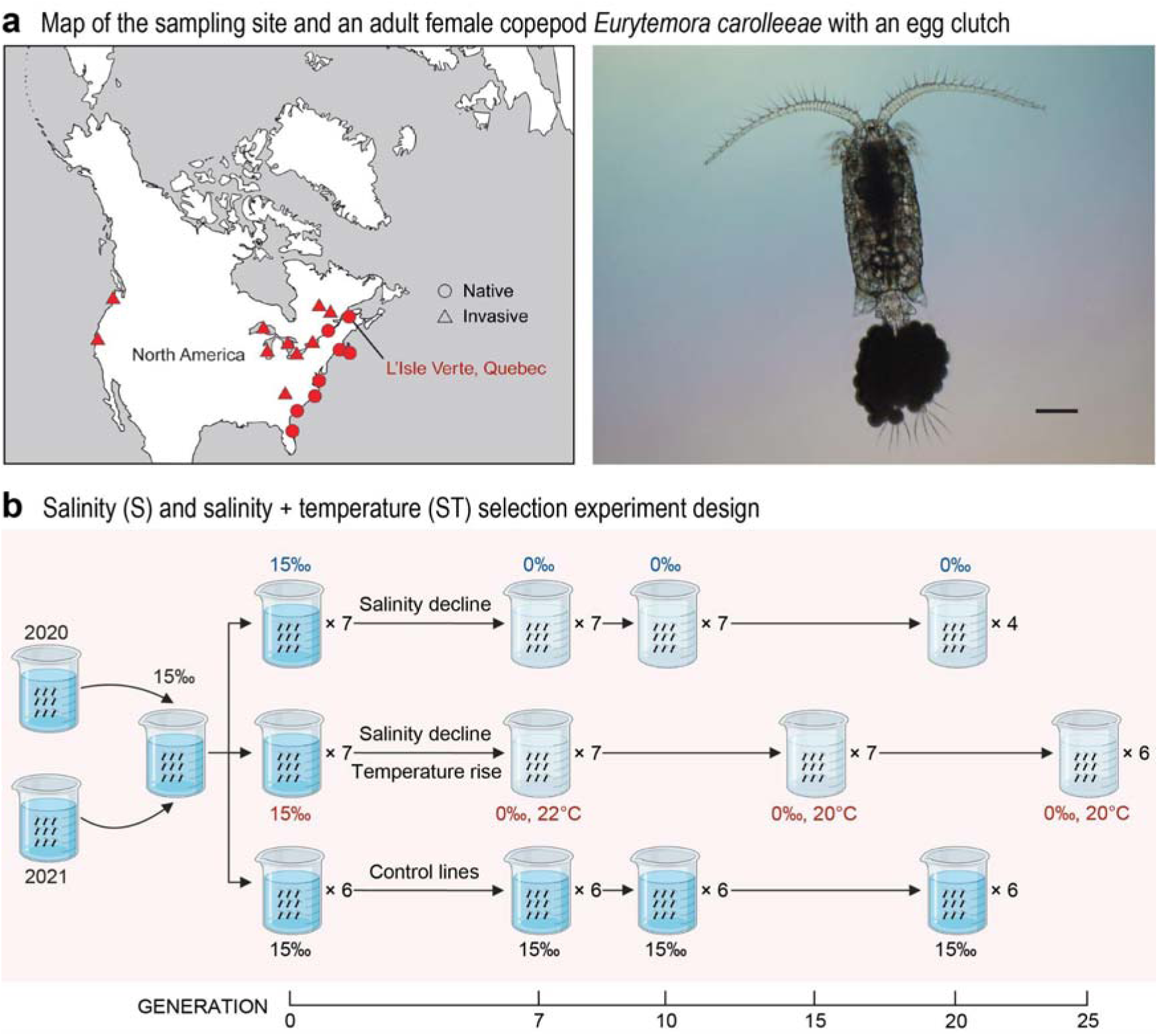
Design of the evolution experiment measuring responses to salinity decline and to salinity decline combined with temperature increase in the copepod Eurytemora carolleeae. **(a)** Map showing the sampling site of the starting population at Baie de L’Isle Verte (St. Lawrence estuary), and a photograph of an adult female E. carolleeae from this location with an egg clutch. Scale bar = 200 μm. Photograph by Violet Krol. **(b)** Design of the Evolve-and-Resequence experiment, using a saline population from the St. Lawrence estuary (see Methods). Top: response to salinity decline only (S). The S selection lines (7 initiated; 4 survived to Generation 20) experienced a stepwise salinity decline from 15‰ to 0‰ over six generations, and were then maintained at 0‰ until Generation 20. Middle: response to salinity decline combined with temperature increase (ST). The ST selection lines (7 initiated; 6 survived to Generation 25) experienced a simultaneous salinity decline (from 15‰ to 0‰ over six generations, then were maintained at 0‰ until Generation 25) and temperature rise (12°C to 22°C over five generations, then maintained at 20°C until Generation 25). Bottom: control lines (6 replicates) remained at a constant salinity of 15‰ and a temperature of 12°C for 20 generations. Numbers next to beakers indicate the number of replicate lines at each stage. Generations 2 to 7 were sampled in all selection and control lines but are not shown in this figure.

This study provides critical insights into the speed, extent, and repeatability of polygenic adaptation when an additional stressor is added to a selection regime. Specifically, here we examine how these adaptive responses are altered when a temperature stressor is added to the well-characterized genomic response to salinity decline in the *E. affinis* complex (Stern & Lee, 2020; Stern et al., 2022; Du et al., 2026). Effects of these two particular stressors, salinity and temperature, are especially relevant for understanding climate change impacts in high-latitude coastal habitats. Impacts of these stressors on *E. affinis* complex populations are particularly important, given the crucial role this copepod plays in supporting several major fisheries (Winkler et al., 2003; Kimmel et al., 2006; Livdāne et al., 2016). In addition, we link the added stressor to fitness consequences and Evolutionary Rescue, with implications for the demography and extinction probability of natural populations. Our results indicate that forecasts based on single-stressor responses may overestimate both the pace and repeatability of adaptation under real-world climate change scenarios.

## Results and Discussion

### Warming impedes the speed and extent of Evolutionary Rescue during salinity adaptation

Evolutionary Rescue occurs when adaptive evolutionary change reverses the demographic decline of a population heading toward extinction, by restoring positive growth through genetic adaptation before extinction occurs (Gomulkiewicz & Holt, 1995; Bell & Gonzalez, 2009; Bell, 2017). Whether a population is rescued depends on multiple factors, including the standing genetic variation available for selection, the rate of environmental change relative to adaptive capacity, and the demographic resilience of the population during the initial fitness decline (Orr & Unckless, 2008; Bell, 2017; Gulisija, 2026).

In the present study, we found that imposing an additional temperature stressor on the salinity selection lines profoundly slowed the speed and hindered the completeness of Evolutionary Rescue (Fig. 2). We subjected replicate selection lines of *E. carolleeae* to two selection regimes, namely salinity decline alone (S lines; 7 replicate lines initiated) versus salinity decline combined with temperature increase (ST lines; 7 replicate lines initiated) (Fig. 1b). We quantified the speed and extent of Evolutionary Rescue in both the salinity alone and salinity + temperature selection regimes by tracking egg production per gravid female across the experiment, using reproductive output as a fitness-related proxy for population recovery. We found that the salinity decline lines (S lines) achieved full Evolutionary Rescue during the experiment while the salinity + temperature lines (ST lines) did not. Specifically, the S lines achieved 103% of the ancestral mean fecundity by Generation 10, whereas the ST lines reached only 68% of the ancestral mean by Generation 17, despite undergoing seven additional generations of selection (Fig. 2).

**Fig. 2.**
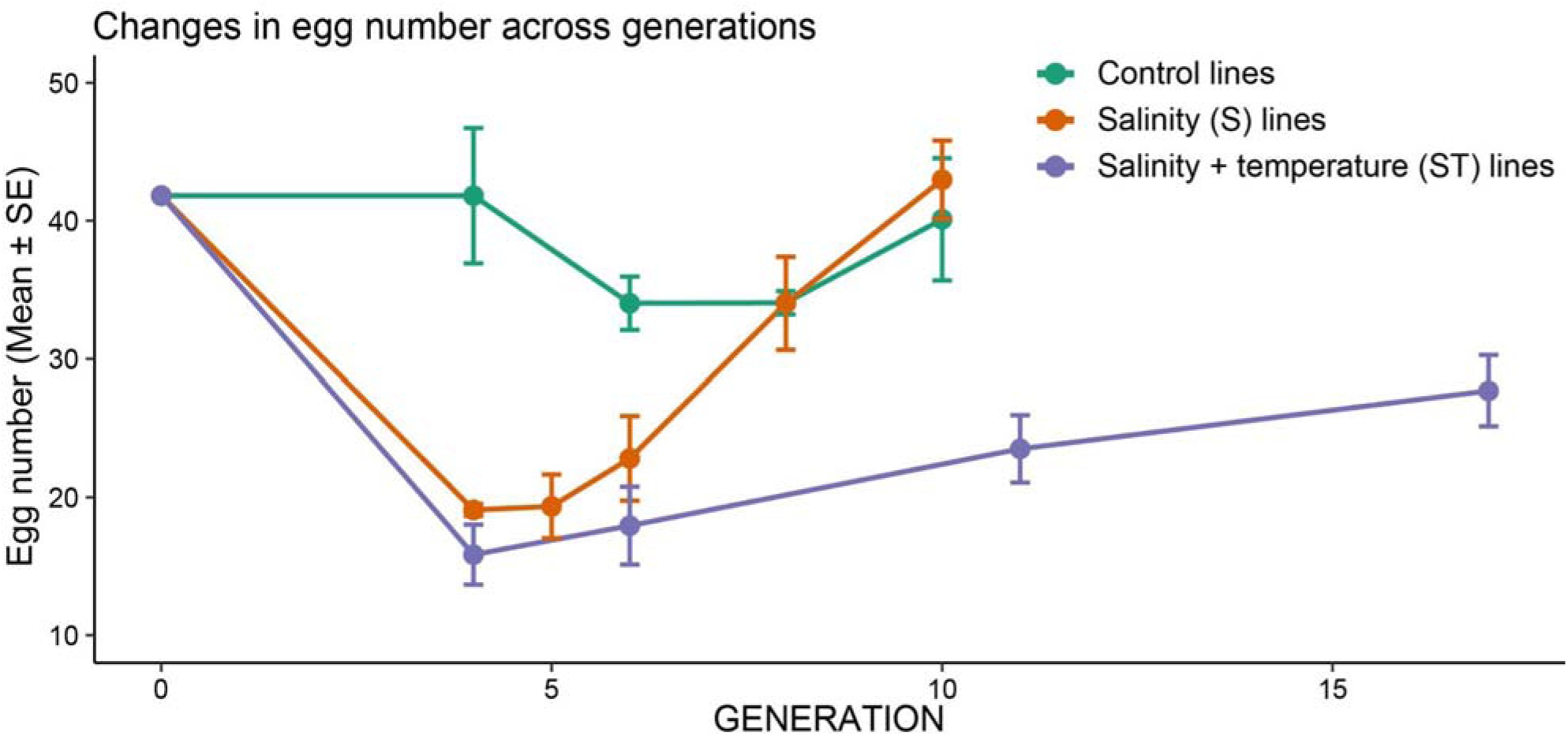
Impact of adding temperature increase to salinity decline on the speed and extent of Evolutionary Rescue. Changes in fitness, measured as mean egg number per gravid female ± SE, during salinity decline alone (S lines, orange) and salinity decline combined with temperature increase (ST lines, purple). Green data points indicate egg numbers of the control lines, in which salinity and temperature remained constant throughout the experiment. The S lines showed an initial reduction in egg number during the salinity decline in the first four generations (from 41.8 ± 2.1 SE to 19.1 ± 0.7 SE eggs), but recovered to ancestral levels by Generation 10 (43.0 ± 1.3 SE eggs per female; N = 70 females across replicate lines). In contrast, the ST lines experienced a steeper initial decline (from 41.8 ± 2.1 SE to 15.8 ± 0.8 SE eggs) and then remained below ancestral and control levels at Generation 17 (28.5 ± 1.2 SE eggs per female; N = 50 females across replicate lines). These results indicate rapid and complete Evolutionary Rescue under salinity decline alone, but delayed and incomplete Evolutionary Rescue when temperature increase was added.

In particular, fitness (in terms of egg number) in the S lines showed a sharp drop at Generation 4, to approximately 46% of the starting value, followed by complete recovery by Generation 10, consistent with rapid and successful Evolutionary Rescue (Fig. 2). Specifically, following the stepwise salinity transition from 15‰ to 0‰, mean fecundity in the S lines dropped sharply from the ancestral baseline of 41.8 ± 2.1 SE eggs per female to 19.1 ± 0.7 SE eggs at Generation 4 (N = 70 females across seven S lines) (Welch’s *t* = 10.53, df = 72, *P* = 3.0 × 10^−16^). The ancestral baseline was estimated from the control lines at Generation 4 (N = 60 females across six control lines; Fig. 2). Egg number remained relatively low at Generation 5 (19.3 ± 1.0 eggs; N = 70; *P* = 7.5 × 10^−16^ versus ancestral), then rose rapidly to 22.8 ± 1.2 SE by Generation 6, 34.0 ± 1.5 SE by Generation 8, and 43.0 ± 1.3 SE by Generation 10 (N = 70 throughout). By Generation 10, egg numbers in the S lines were statistically comparable to the ancestral baseline (41.8 ± 2.1 SE eggs; *t* = 0.48, df = 101, *P* = 0.63) and to contemporaneous control lines (40.1 ± 1.8 SE eggs; *t* = 1.31, df = 110, *P* = 0.19).

In contrast, adding temperature increase to the salinity decline selection regime (ST lines) produced a greater initial fitness decline, to approximately 38% of the starting value, followed by slower recovery and incomplete restoration of fitness within the experimental timeframe (Fig. 2). The ST lines reached a significantly deeper fitness trough of 15.8 ± 0.8 SE eggs at Generation 4 (N = 70 females across seven ST lines; 3.3 eggs lower than the matched S time point; Welch’s *t* = 3.03, df = 134, *P* = 3.0 × 10^−3^) and recovered much more slowly than the S lines. Fecundity in the ST lines rose to only 17.9 ± 1.2 SE eggs by Generation 6 (N = 70), 23.5 ± 1.3 SE eggs by Generation 11 (N = 35), and 28.5 ± 1.2 SE eggs by Generation 17 (N = 50). Although this increase represented a significant recovery relative to the ST lines at Generation 4 (*t* = 8.69, df = 91, *P* = 1.3 × 10^−13^), the egg number of 28.5 ± 1.2 SE in the ST lines at Generation 17 remained significantly lower than the ancestral baseline (41.8 ± 2.1 SE eggs; *t* = 5.62, df = 93, *P* = 1.9 × 10^−7^), the control lines at Generation 10 (40.1 ± 1.8 SE eggs; *t* = 5.41, df = 100, *P* = 4.3 × 10^−7^), and the S lines at Generation 10 (43.0 ± 1.3 SE; *t* = 8.29, df = 117, *P* = 2.2 × 10^−13^).

The delayed and incomplete fitness recovery when warming was added to the selection regime aligns with several theoretical predictions regarding constraints on Evolutionary Rescue. Fisher’s geometric model, extended to populations facing simultaneous selection on multiple physiological axes, predicts a “cost of complexity” whereby the rate of adaptation declines as the dimensionality of selection increases. This cost arises because mutations that simultaneously improve all selected traits become rarer as the number of traits grows (Orr, 2000; Tenaillon, 2014). Lindsey et al. (2013) further demonstrated that the probability of Evolutionary Rescue falls steeply as the rate of environmental change accelerates, because populations have less time to mount an adaptive response before extinction. From the perspective of any single physiological trait, adding temperature stress in addition to salinity stress effectively increases the severity and dimensionality of environmental change. Consistent with the critical rate of environmental change beyond which extinction becomes inevitable for a given trait architecture and population size (Bürger & Lynch, 1995), our results suggest that adding a second concurrent stressor moved populations closer to this boundary, although not beyond it. As a result, the ST lines persisted but only partially recovered during the timeframe of our experiment.

### Warming alters the genome-wide polygenic selection response to salinity decline

Both the S and ST selection regimes elicited genome-wide polygenic responses, but adding temperature increase substantially altered the underlying genomic response to selection (Figs. 3 and 4; Table S1). Adding warming produced a more polygenic response, recruiting approximately 50% more candidate SNPs than salinity decline alone (Fig. 3; Table S1). It also produced a weaker per-locus selection response, with a slower rise in frequencies of beneficial alleles (Fig. 4b versus 4a) and a 35% reduction in the mean per-locus selection coefficient (Fig. 4f versus 4e).

**Fig. 3.**
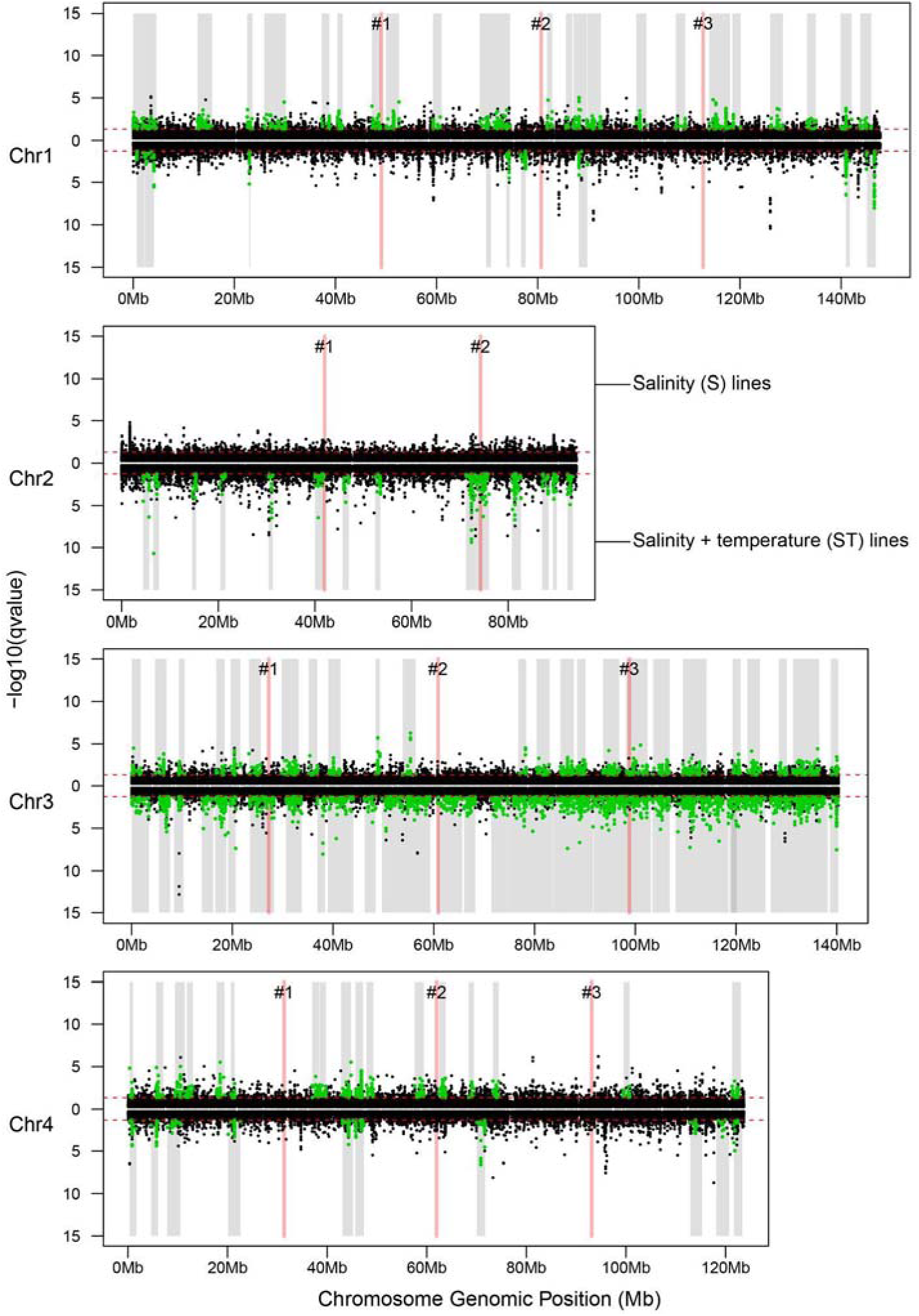
Manhattan plots of genomic signatures of selection across all four chromosomes of the Eurytemora carolleeae genome. For each chromosome (Chr1–Chr4), the upper portions display –log10(q-value) from the piecewise linear mixed model (LMM) for the salinity (S) selection lines, whereas the lower portions show the corresponding LMM results for the salinity + temperature (ST) selection lines. Red dashed horizontal lines indicate the significance threshold (q < 0.05). Green dots denote SNPs that exceeded the significance threshold and fell within identified haplotype blocks. Black dots represent background genomic SNPs. Gray shaded bars delineate the boundaries of haplotype blocks identified as targets of selection on each chromosome. Numbered pink vertical bars (#1, #2, #3) mark the positions of chromosomal fusion sites previously identified in the Atlantic clade genome (Du et al., 2025). Selection targets were distributed genome-wide in both experiments but showed differences in genomic distribution. Notably, Chromosome 2 harbored specific selection signatures in the ST lines, but not in the S lines.

**Fig. 4.**
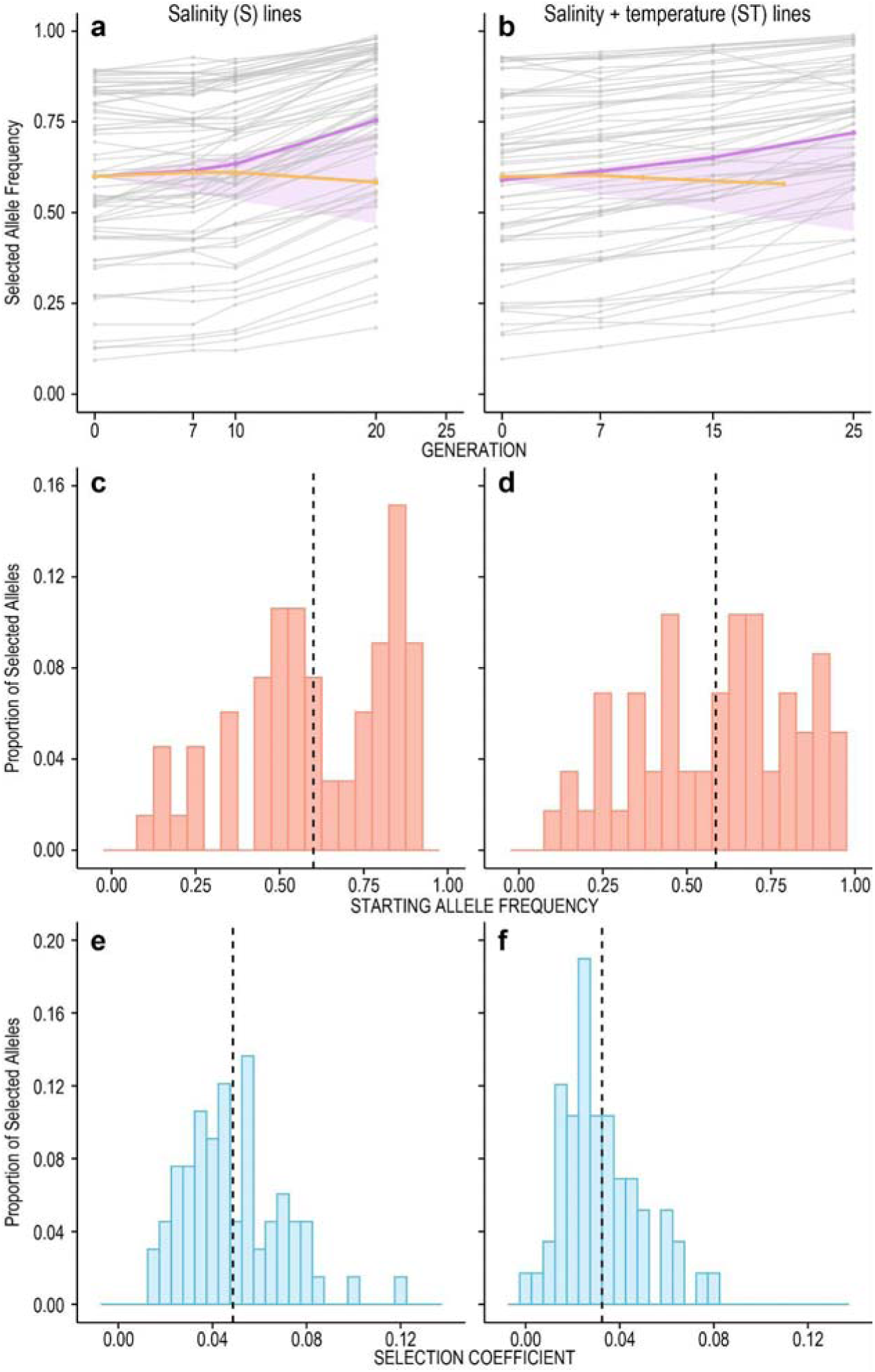
Contrasting selection responses in the salinity decline versus salinity decline + warming evolution experiments of Eurytemora carolleeae. (a, **b)** Allele frequency trajectories of selected haplotype blocks during **(a)** the salinity decline (S) experiment (2,796 SNPs in 66 haplotype blocks) and **(b)** the salinity decline + temperature increase (ST) experiment (4,362 SNPs in 58 haplotype blocks). Gray lines represent the mean allele frequency of each selected haplotype block across surviving replicate selection lines (N = 4 surviving lines in the S experiment; N = 6 surviving lines in the ST experiment). Purple lines represent the mean frequency of all selected haplotype blocks in the selection lines. Orange lines denote the corresponding mean in the control lines. Pink shaded areas indicate the 1st and 99th percentile ranges from 10,000 neutral drift simulations. **(c, d)** Starting allele frequency distributions for selected haplotype blocks in the **(c)** S and **(d)** ST experiments. For each selected haplotype block, allele (haplotype block) frequencies were polarized to the allele that increased in frequency during selection, rather than to ancestral or derived allelic states. Thus, the plotted starting frequency represents the initial frequency of the putatively favored allele in the ancestral population. Dashed vertical lines indicate mean starting allele frequencies. **(e, f)** Distributions of estimated selection coefficients for selected haplotype blocks in the **(e)** S and **(f)** ST experiments. Dashed vertical lines indicate the mean selection coefficients.

Specifically, adding warming to the salinity decline regime recruited approximately 1.50-fold as many candidate SNPs, from 10,176 to 15,272 candidate SNPs, into the selection response as salinity decline alone (Table S1). To identify the SNPs under selection, we applied Cochran–Mantel–Haenszel (CMH) tests, Chi-square tests, and a piecewise linear mixed model (LMM) to the time-resolved pooled whole-genome sequencing data sampled across generations, up to Generation 20 for the S lines and Generation 25 for the ST lines (see Methods for details). From the 168,532 biallelic SNPs anchored to the four *E. carolleeae* chromosomes, the S lines yielded 10,176 candidate SNPs with signatures of selection (combined non-redundant set from the three tests; CMH: 799; Chi-square: 1,459; LMM: 9,410), whereas the ST lines yielded 15,272 candidate SNPs (CMH: 1,968; Chi-square: 2,322; LMM: 13,242) (Figs. S1 to S8; Table S1).

Notably, for the piecewise structure of the LMM analysis, the time periods before and after Generation 7 were analyzed separately, corresponding to the end of the salinity and temperature transition. Separate analyses were necessary because allele frequency trajectories differed between the early transition phase, when the environmental variables were changing, and the later phase, when the selection lines were held under the final constant conditions. As such, a single linear model across the full trajectory would likely underestimate or obscure the allele frequency changes during the early transition phase. The piecewise structure also accommodated the mismatched sampling time points between the S (Generations 10 and 20) and ST (Generations 15 and 25) lines, which arose from differences in developmental rate of the copepods at different temperatures (see Methods).

The larger number of candidate SNPs recruited under the salinity + temperature regime (ST lines) was consistent with salinity and temperature acting largely on distinct genomic targets under selection. Thus, adding warming enlarged the pool of loci available to respond to selection. This result was consistent with previous findings in wild *E. affinis* proper populations, where distinct sets of SNPs were associated with salinity and temperature gradients with relatively low overlap (Diaz et al., 2026).

Moreover, we found that adding warming to the selection regime resulted in striking shifts in the genomic distribution of the selection response, with pronounced changes in the concentration and localization of selected haplotype blocks on the different chromosomes (χ² = 21.82, df = 3, *P* = 7.1 × 10^−5^; Table S1). To account for genetic linkage among nearby SNPs, candidate SNPs were grouped into selected haplotype blocks using the haplovalidate approach (Otte & Schlötterer, 2021b). The S lines yielded 66 haplotype blocks containing 2,796 SNPs in total, whereas the ST lines yielded 58 haplotype blocks containing 4,362 SNPs (Figs. 3 and 4a, b; Figs. S1 to S8; Tables S1 to S3). Notably, the chromosomal distribution of these blocks differed sharply between the two selection regimes. Under salinity decline alone, the selected blocks were concentrated on Chromosomes 1, 3, and 4 (25, 24, and 17 blocks, respectively; 37.9%, 36.4%, and 25.8% of the 66 S blocks), with zero selected blocks on Chromosome 2. When warming was added to the selection regime, selected blocks were distributed across all four chromosomes (10, 14, 24, and 10 blocks on Chromosomes 1 to 4, respectively; 17.2%, 24.1%, 41.4%, and 17.2% of the 58 ST blocks). Chromosome 2 alone harbored 14 blocks spanning 19.26 Mb of selected genomic regions in the ST lines, none of which were detected under salinity decline alone (Fig. 3; Tables S1 and S4). On the other hand, the ST lines exhibited 15 fewer selected haplotype blocks than the S lines on Chromosome 1 and 7 fewer haplotype blocks on Chromosome 4 (Fig. 3; Table S1). Thus, adding warming did not simply add temperature-responsive loci on top of the salinity response within the same selected blocks. Instead, the ST lines both recruited otherwise unselected chromosomal regions and also lost some of the selected haplotype blocks present in the S lines.

Despite the lower number of haplotype blocks in the ST lines, the ST blocks were on average larger than the S blocks (median size 1.82 Mb for ST blocks versus 1.76 Mb for S blocks) and had a wider range of values (0.07 to 11.38 Mb for ST blocks versus 0.68 to 5.20 Mb for S blocks). In addition, ST blocks contained more SNPs per block (75.2 versus 42.4 on average). This result was consistent with selection acting on larger linked chromosomal segments in the ST lines when multiple physiological traits were under simultaneous selection.

A striking quantitative difference between the two selection regimes was the 35% reduction in the mean per-locus selection coefficient (*s*) in the ST lines, relative to the S lines (S lines: *s* = 0.049 ± 0.003 SE, range 0.013 to 0.12; ST lines: *s* = 0.032 ± 0.002 SE, range 0 to 0.078; Mann–Whitney *U* = 2,816, *P* = 6.3 × 10^−6^; Fig. 4e, f; Tables S2 and S3). The genome-wide Spearman rank correlation in the selection coefficients between the two regimes was relatively low (ρ = 0.265, N = 168,532 SNPs, *P* < 2.2 × 10^−16^). The selection response was weaker in the ST lines even though selection under both regimes acted on similar levels of standing variation, with similar starting mean frequencies of beneficial alleles (S: 0.60 ± 0.028 SE; ST: 0.59 ± 0.031 SE; Mann–Whitney *U* = 1,987, *P* = 0.72; Fig. 4c, d; Tables S2 and S3). That is, in both selection regimes, selection acted on pre-existing intermediate-frequency variants rather than on *de novo* mutations or rare alleles.

These results indicate that the per-locus rate at which intermediate-frequency alleles were driven toward fixation differed substantially between the two selection regimes. A higher mean selection coefficient corresponds to a faster per-generation increase in adaptive allele frequency and a greater individual-locus contribution to fitness recovery within a given number of generations. In contrast, a lower mean selection coefficient implies that each locus tends to contribute less toward adaptation per generation, so that more loci with smaller individual effects are needed to achieve a comparable phenotypic shift. Consistent with this contrast, individual selection coefficients per locus reached up to *s* = 0.12 in the S lines (Table S2) but did not exceed *s* = 0.078 in the ST lines (Table S3). In addition, the full distribution of *s* shifted toward lower values when warming was added, with ST blocks over-represented in the *s* = 0.01 to 0.03 class and depleted at *s* > 0.05 (Fig. 4e, f). Notably, the time required for an adaptive allele to rise in frequency, and therefore the time for a polygenic response to restore fitness, scales inversely with *s* (Hermisson & Pennings, 2005; Höllinger et al., 2019). Thus, the lower selection coefficients under added warming provide a direct mechanistic link to the slower and incomplete fitness recovery observed in our selection lines (Fig. 2). This simultaneous broadening (more loci) and shallowing (lower per-locus *s*) of the polygenic response in the ST selection regime is the quantitative signature predicted by theory for selection on a higher-dimensional fitness landscape (Orr, 2000; Pritchard & Di Rienzo, 2010; Tenaillon, 2014).

### Warming reduces parallelism among replicate selection lines during salinity decline

Adding temperature increase to the salinity decline regime greatly reduced parallelism among replicate selection lines, relative to salinity decline alone (Fig. 5). Because low parallelism can arise either from limited sharing of selected loci or from differences in the direction and magnitude of allele frequency changes across selection lines, we quantified parallelism using two complementary metrics. Specifically, we examined the mean pairwise Jaccard index, which measures the proportion of selected haplotype blocks shared between pairs of replicate lines at a given generation, capturing whether the same loci respond to selection across the replicate lines. On the other hand, we also performed principal component analysis (PCA) of allele frequency changes at selected SNPs to capture the direction and magnitude of the multivariate genomic response. In addition to our definition of “parallelism” stated in the Introduction, we further define this term here to refer to the sharing of selected haplotype blocks among replicate lines. Parallelism defined here is distinct from convergence, which would additionally require the variance among replicate lines to decline over time (Bolnick et al., 2018). Both the Jaccard index and PCA results indicated that adding warming to the salinity decline selection regime reduced the extent of parallelism among the selection lines.

**Fig. 5.**
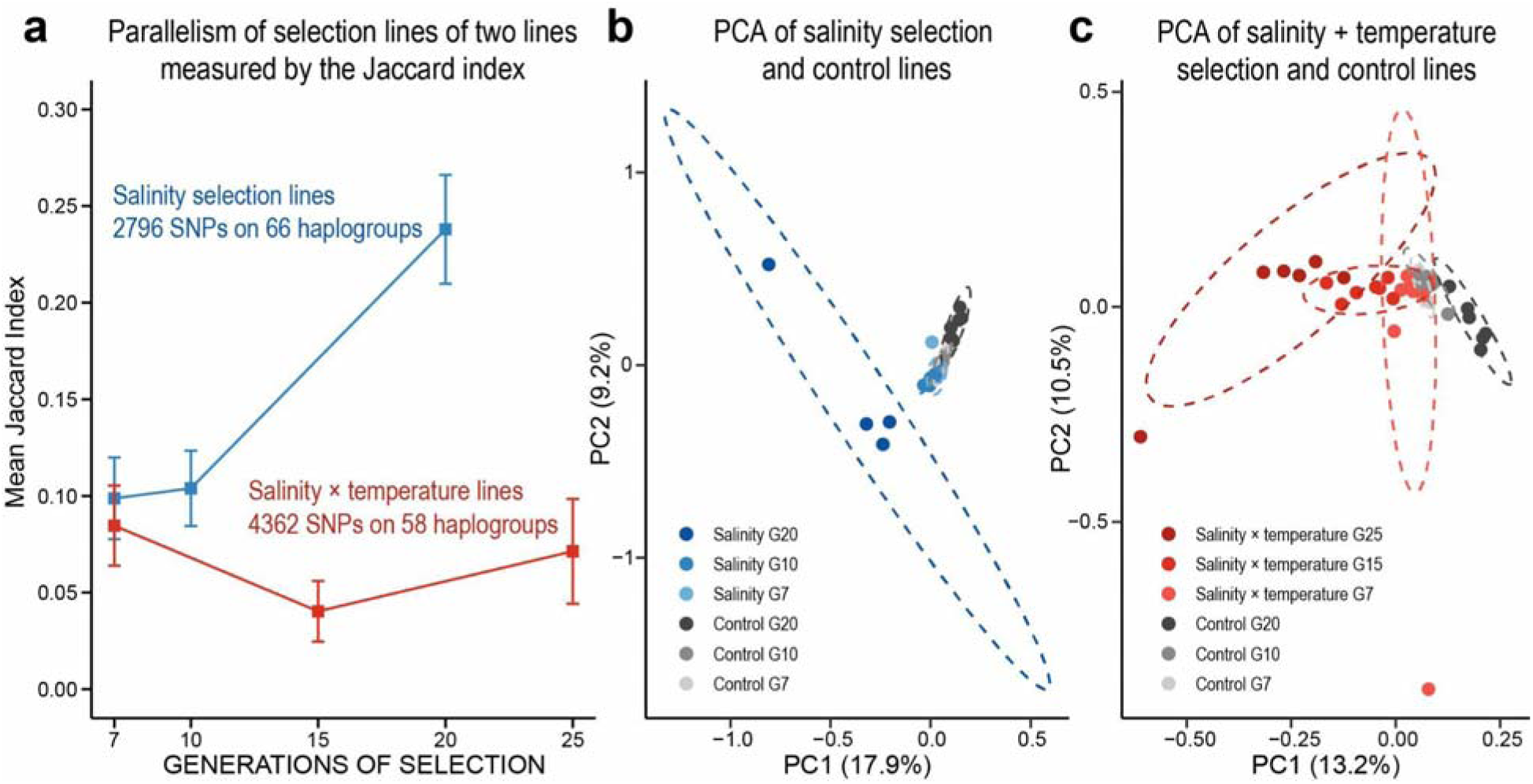
Parallelism in allele frequency shifts among replicate selection lines of Eurytemora carolleeae under salinity decline alone (S lines) versus salinity decline combined with temperature increase (ST lines). **(a)** Mean pairwise Jaccard index measuring the proportion of selected haplotype blocks shared between pairs of replicate lines, plotted across generations for the S lines (blue; based on 2,796 SNPs across 66 haplotype blocks) and ST lines (red; based on 4,362 SNPs across 58 haplotype blocks). Data points are the mean ± SE across all pairwise comparisons among replicate lines at each generation. Parallelism among the S lines rose from 0.099 ± 0.021 SE at Generation 7 to 0.238 ± 0.028 SE by Generation 20. In contrast, parallelism among the ST lines remained near 0.08 throughout the experiment, reaching only 0.071 ± 0.027 SE at Generation 25 (Mann–Whitney U = 78, P = 0.0084 comparing terminal-generation pairwise Jaccard distributions). Pairwise Jaccard index values for all generations are provided in Tables S5 and S6. **(b)** Principal component analysis (PCA) of allele frequency profiles at selected SNPs in the S experiment. By Generation 20, three of the four surviving S lines clustered together at negative PC1 and PC2 values, while the fourth surviving line was an outlier, well separated from the control lines. **(c)** PCA of allele frequency profiles in the ST experiment. ST lines at Generations 7 and 15 remained proximate to the control lines, with an extreme outlier at Generation 7. By Generation 25, the six surviving ST lines had separated from the control lines along PC1 with one extreme outlier; however, these ST lines remained more dispersed from one another than the three clustered S lines at Generation 20 (b).

Based on the Jaccard index, by the end of the evolution experiments, parallelism among replicate lines was far lower when warming was added (ST lines) than under salinity decline alone (S lines). The mean pairwise Jaccard index reached 0.238 ± 0.028 SE in the S lines at Generation 20, but only reached 0.071 ± 0.027 SE in the ST lines at Generation 25, despite five additional generations of selection (Mann–Whitney *U* = 78, *P* = 0.0084; Fig. 5a; Tables S5 and S6). Parallelism among the ST lines after 25 generations (0.071 ± 0.027 SE) was even lower than among the S lines after only seven generations of selection (0.099 ± 0.021 SE). This low parallelism was consistent with the broader (more loci) and weaker selection response (lower per-locus *s*) of the selected loci involved (see previous section). That is, the selection response to salinity + temperature change (ST lines) recruited roughly 50% more candidate SNPs with a 35% lower mean per-locus selection coefficient than the response to salinity decline alone (S lines) (Fig. 4; Table S1). As selection in the ST lines was distributed across more loci of smaller effect, with a greater fraction of those loci potentially becoming functionally redundant, different replicate lines could recruit different subsets of these loci during adaptation. The probability that any two lines independently arrive at the same subset of selection alleles therefore becomes low (Barghi et al., 2019; Láruson et al., 2020; Du et al., 2026).

In particular, parallelism among the S lines increased over the course of the experiment, from a mean pairwise Jaccard index of 0.099 ± 0.021 SE at Generation 7 (21 pairwise comparisons among 7 lines), to 0.104 ± 0.020 SE at Generation 10 (21 pairs), and finally to 0.238 ± 0.028 SE by Generation 20 (6 pairs among 4 surviving lines; Fig. 5a; Table S5). On average, each S line at Generation 20 possessed 22.25 ± 6.34 SE (33.7%) of the 66 selected blocks. This 2.3-fold rise in the pairwise Jaccard index from Generation 10 to Generation 20 indicates that replicate S lines drew on an increasingly shared set of selected haplotype blocks as adaptation proceeded. This pattern was consistent with a repeatable selection response built from a limited set of relatively large-effect salinity-adaptive alleles (haplotype blocks) present at intermediate frequencies in the founder population (mean starting frequency = 0.60 ± 0.028 SE; Fig. 4c).

In sharp contrast, the ST lines did not achieve comparable levels of parallelism at any sampled generation, with the mean pairwise Jaccard index remaining near 0.08 throughout the experiment. Specifically, the Jaccard index remained low across generations, at 0.085 ± 0.021 SE at Generation 7 (21 pairs among 7 lines), then 0.040 ± 0.016 SE at Generation 15 (21 pairs), and finally 0.071 ± 0.027 SE at Generation 25 (15 pairs among 6 surviving lines; Fig. 5a; Table S6). On average, each ST line at Generation 25 possessed only 5.67 ± 2.74 SE (9.8%) of the 58 selected blocks.

Principal component analysis (PCA) of allele frequency changes at selected SNPs corroborated the contrasting patterns of parallelism between the S and ST lines (Fig. 5b, c). In the S experiment, selection lines at Generations 7 and 10 clustered tightly with control lines in PCA space (within-group mean centroid distance ≤ 0.05) (Fig. 5b). By Generation 20, the four surviving S lines separated from the control lines along the PC1 axis (17.9% of variance explained; S mean PC1 = –0.39 ± 0.14 SE; control mean PC1 = 0.13 ± 0.01 SE). Three of these four S lines clustered together, while the fourth line was an outlier. Despite the limited number of lines surviving to Generation 20, the clear separation of S lines from control lines and the tight clustering of three of the four S lines were consistent with the moderate selection response and the rising parallelism indicated by the Jaccard index (Fig. 5a, b).

In contrast, the ST lines diverged more gradually from the control lines and reached a lower magnitude of change in the PCA space relative to S lines (Fig. 5c). Specifically, the ST lines at Generations 7 and 15 remained proximate to the control lines along PC1 (13.2% of variance explained) and PC2 (10.5% of variance explained), except for an extreme outlier at Generation 7. At Generation 15, the ST lines clustered close to one another and to the control lines (within-group centroid distance = 0.05). By Generation 25, the six surviving ST lines separated from the control lines along PC1 with one extreme outlier (Fig. 5c) (ST mean PC1 = – 0.29 ± 0.07 SE; control mean PC1 = 0.18 ± 0.01 SE), but with a smaller absolute multivariate shift than the S lines at Generation 20 (within-group centroid distance = 0.16 ± 0.06 SE versus 0.39 ± 0.26 SE). This smaller shift was consistent with the lower per-locus allele frequency shifts in the ST lines relative to S lines, implied by the lower selection coefficients (mean *s* = 0.032 versus 0.049). In addition, the lower separation of the ST lines from the control lines was consistent with the weak selection response in the ST lines (Fig. 5c). This weak selection response, along with other factors (see section below on *Mechanisms underlying the reduced speed and parallelism under added warming stress*), contributed to the low levels of parallelism in the ST lines throughout the experiment, as indicated by the low Jaccard scores (Fig. 5a).

### Warming recruits divergent genomic targets of selection under salinity decline

Our results revealed that adding temperature increase to the selection regime substantially recruited divergent genomic targets under selection among replicate selection lines relative to salinity decline alone (Fig. 3). We assessed the extent to which the salinity versus salinity + temperature selection involved the same genetic targets at multiple levels of genomic organization. That is, we compared the targets of selection between the S and ST lines at the levels of haplotype blocks, individual SNPs, and genes. In addition, we compared the functional categories of genes under selection based on Gene Ontology (GO) enrichment in the S versus ST experiments.

At the level of physical genomic regions (haplotype blocks), we found a total of 68.2 Mb of haplotype blocks shared between the S and ST selection regimes (Jaccard index = 0.30; Table S4). This sharing was driven mostly by Chromosome 3, where the per-chromosome Jaccard index reached 0.45. Sharing of haplotype blocks was intermediate on Chromosome 1 (Jaccard index = 0.15) and Chromosome 4 (Jaccard index = 0.21) and absent on Chromosome 2 (Table S4). On the other hand, at the level of individual candidate SNPs, the overlap was much lower between the selection regimes. We found only 3,174 of the 22,274 unique candidate SNPs shared between the two selection regimes (Jaccard index = 0.14; Fig. S9a). When restricting the comparison to only the SNPs contained within selected haplotype blocks in both regimes, we found only 549 SNPs shared between the regimes (Jaccard index = 0.083; Fig. S9b), representing 19.6% of the selected haplotype block SNPs in the S lines and 12.6% in the ST lines.

For the selected genes containing selected SNPs within the haplotype blocks, we found 306 genes shared between the two selection regimes (Tables S7 to S9). These 306 shared genes represented 39.9% of the 766 genes underlying selected blocks in the S lines (Table S7) and 27.7% of the 1,106 genes in the ST lines (Table S8) (Jaccard index = 0.20; Table S10). This gene-level overlap was higher than SNP-level overlap (Jaccard index = 0.14), as expected, because a single gene can contain different selected SNPs in the two selection regimes. The low overlap in shared SNPs between the S and ST experiments was somewhat consistent with the wild-population observation that only 5 to 18% of significant SNP associations were shared between salinity and temperature responses across *E. affinis* proper populations (Spearman correlations between the two variables of only ρ = 0.07 to 0.19) (Diaz et al., 2026).

We also investigated how the two selection regimes differed in the functional pathways involved by performing GO enrichment analysis of the genes containing selected haplotype block SNPs (Tables S11 and S12). The S regime yielded 230 significantly enriched GO terms, and the ST regime yielded 260, but only 33 terms were shared (14.3% of S terms and 12.7% of ST terms; Jaccard index = 0.072; Tables S10 and S13). The shared terms were mostly related to neurological processes, including calcium ion transmembrane import (GO:0097553), regulation of neurotransmitter levels (GO:0001505), and nervous system process (GO:0050877) (Table S13). Notably, the overlap in functional categories (14.3% of S terms) was even lower than the overlap in the underlying gene targets (39.9% of S genes), indicating that shared genes were often assigned to different enriched functions in the two regimes (Table S10).

Under salinity decline alone, enriched GO categories were dominated by ion channel functions, neurological processes, and membrane regulation, including chloride channel activity (GO:0005254), transmitter-gated monoatomic ion channel activity (GO:0022824), ligand-gated ion channel signaling pathway (GO:1990806), regulation of membrane potential (GO:0042391), and chaperone binding (GO:0051087) (Table S11). Several of these categories matched the neurological and ion regulatory processes known to underlie freshwater adaptation from prior laboratory evolution experiments (Stern et al., 2022; Du et al., 2026) and wild population genomic surveys in the Atlantic and Europe clades of the *E. affinis* complex (Lee, 2023; Du et al., 2025; Diaz et al., 2026).

When warming was added to the salinity decline selection regime, the enriched categories broadened and shifted. The most notable shift was from the enrichment of ion channels and membrane regulation under salinity decline alone toward the enrichment of ion transport, ion homeostasis, structural, and metabolic machinery in the ST lines. The significantly enriched ion regulatory GO terms in the ST lines included monoatomic cation transmembrane transporter activity (GO:0008324), inorganic cation transmembrane transporter activity (GO:0022890), intracellular monoatomic ion homeostasis (GO:0006873), and epithelial fluid transport (GO:0042045) (Table S12). Beyond ionic regulation, the ST selection regime additionally recruited categories such as cytoskeletal protein binding (GO:0008092), respiratory system process (GO:0003016), ATP hydrolysis activity (GO:0016887), and hormone metabolic process (GO:0042445). These added functional categories in the ST lines were consistent with thermal stress in ectotherms imposing selection on membrane fluidity, protein stability, mitochondrial function, and metabolic rate that salinity decline alone would not impose (Angilletta et al., 2002; Hoffmann & Sgrò, 2011).

The even greater involvement of ion transport-related genes in the ST lines relative to the S lines was intriguing. This result suggests that salinity adaptation under warming required even greater selection on ion transport-related genes than under salinity decline alone. In our prior evolution experiments, ion transport-related functions comprised the top GO categories under selection in response to salinity decline in *E. affinis* complex populations (Stern et al., 2022; Du et al., 2025, 2026). In addition, our previous comparative studies revealed the evolution of ion transporter expression and activity between ancestral saline and freshwater invading populations of the *E. affinis* complex (Posavi et al., 2020; Popp et al., 2024; Lee et al., 2011). In wild *E. affinis* complex populations, ion transport-related functions were significantly enriched along the salinity gradient (Stern & Lee, 2020; Diaz et al., 2026), but not along the temperature gradient (Diaz et al., 2026). This result suggests that temperature change alone would not induce the involvement of ion transport-related functions. Our results here suggest that temperature increase likely magnifies the effects of salinity stress and induces a greater salinity selection response in *E. affinis* complex populations. Additional studies that include additional temperature stress treatments would be needed to determine the impacts of temperature stress on salinity adaptation.

### Mechanisms underlying the reduced speed and parallelism under added warming stress

The slower fitness recovery and lower extent of parallelism among replicate lines under salinity + temperature selection, relative to salinity decline alone, likely reflected the effects of several reinforcing mechanisms. These include increases in the dimensionality of the fitness landscape (“cost of complexity”) and an increased redundancy of the more polygenic response under the salinity + temperature selection regime. An additional mechanism includes the linkage of alleles due to the fused genome architecture of *E. carolleeae* (N = 4 chromosomes). Linkage due to fewer chromosomes would constrain how selection can act on physically linked salinity- and temperature-responsive alleles. All of these factors might be contributing to the slower and incomplete fitness recovery of the ST lines (Fig. 2).

Adding temperature stress to salinity decline appears to have increased the dimensionality of the fitness landscape, imposing what Orr (2000) termed the “cost of complexity.” As more phenotypic axes must improve simultaneously in response to selection, the proportion of genetic changes that are beneficial across all axes declines, and the per-generation rate of adaptation is expected to decrease (Tenaillon, 2014). Consistent with salinity and temperature acting on largely distinct traits and loci, the genome-wide Spearman rank correlation between the per-SNP selection coefficients of the two regimes was significant but weak (ρ = 0.265 across 168,532 SNPs, *P* < 2.2 × 10^−16^), and the SNP-level overlap (Jaccard index) was only 0.14. This pattern matches our prior genomic survey of wild Baltic Sea and North Sea region populations, which found weak correlations between salinity- and temperature-associated SNPs (ρ = 0.07 to 0.19; Diaz et al., 2026). Because salinity and temperature adaptation largely involve distinct physiological traits and loci, imposing selection on both sets of loci simultaneously likely increases the effective dimensionality of adaptation.

Adding warming to the selection regime is also expected to increase the number of loci that contribute to adaptation, thereby increasing the genetic redundancy of the loci and the number of alternative genetic combinations that contribute to adaptation (Barghi et al., 2019, 2020; Láruson et al., 2020). As the number of loci contributing to adaptation grows, a greater fraction of loci become functionally interchangeable, such that replicate populations can reach comparable phenotypes through different subsets of loci, reducing parallelism. Under salinity decline alone, a core set of ion regulatory and neurological alleles and pathways is repeatedly recovered across laboratory experiments, wild populations, and divergent clades of the *E. affinis* complex (Stern et al., 2022; Lee, 2021, 2023; Posavi et al., 2020; Du et al., 2025, 2026; Diaz et al., 2026). In this study, replicate S lines accordingly arrived at increasingly overlapping sets of selected loci (Fig. 5a). When warming was added to the selection regime, a broader range of physiological functions and genomic regions responded to selection (Tables S8, S10, S12). This increase in contributing loci likely resulted in replicate lines recruiting different allelic combinations, consistent with the persistently low Jaccard index values in the ST lines (Fig. 5a).

The effect of linkage due to the fused genome architecture of *E. carolleeae* likely constrains selection by physically linking salinity- and temperature-responsive alleles within large chromosomal regions. The *E. carolleeae* genome contains only four chromosomes, formed by multiple ancient chromosomal fusions (Du et al., 2025). Having fewer chromosomes reduces the independent assortment of loci on the same chromosome and is consistent with the large selected haplotype blocks observed in this study (median 1.76 Mb in S and 1.82 Mb in ST). When salinity- and temperature-responsive loci lie within the same linked haplotype, the response of that haplotype depends on its joint allelic composition. This linkage would make it difficult for selection to act on the two sets of loci independently (Hill–Robertson effect; Comeron et al., 2008). Different replicate lines may then rise in frequency for alternative multilocus haplotypes that each provide a different partial adaptive solution, further reducing parallelism.

Finally, a direct consequence of the broader genomic response under added warming is the weaker per-locus selection coefficient (mean *s* = 0.032 versus 0.049). Weaker per-locus selection reduces the deterministic component of allele frequency change relative to genetic drift (Höllinger et al., 2019; Schlötterer, 2023), which increases stochastic differences among replicate evolutionary trajectories (Franssen et al., 2017a; Höllinger et al., 2019). Together, the higher dimensionality of selection, the greater genetic redundancy, and the linkage imposed by a highly fused genome are consistent with the more polygenic, weaker, and less parallel response we observed when warming was added to the salinity decline selection regime.

Notably, our experimental design allowed us to attribute the effects of slower fitness recovery and lower among-replicate parallelism to the addition of temperature to our selection regime. However, we did not investigate the effects of temperature alone. Thus, we cannot determine the extent to which the selection response we found in the ST lines was due to the interaction effects between salinity and temperature or due to the effects of temperature alone. We plan to explore this topic further using a three-regime design, including salinity-only, temperature-only, and combined treatments under matched conditions, to fully resolve these effects and their underlying mechanisms.

### Implications for evolutionary forecasting under climate change

Our results have direct implications for forecasting the capacity of natural populations to evolve and persist under realistic multi-stressor impacts due to climate change. The copepod *E. affinis* species complex is an important component of zooplankton communities in many coastal habitats, serving as a major food source for important fisheries across the Northern Hemisphere (Winkler et al., 2003; Kimmel et al., 2006; Livdāne et al., 2016). High-latitude coastal habitats are currently experiencing rapid rates of simultaneous freshening and warming (Durack et al., 2012; Lavoie et al., 2015; Long & Perrie, 2015; Gröger et al., 2019), which have disruptive impacts on coastal food webs and patterns of ocean circulation (Rahmstorf, 2024). Our laboratory experiment revealed that adding warming to the salinity decline regime intensified the fitness cost and produced a genomic response that deviated from the salinity decline-only experiment (Stern et al., 2022; Du et al., 2026). Under combined salinity decline and warming, the slower and incomplete fitness recovery and weaker per-locus selection indicate the much greater challenge of adapting to more than one environmental variable. Such a greater challenge of adapting to two simultaneous stressors has also been found for the copepod *Acartia tonsa* in response to ocean warming and acidification (Dam et al., 2021; Brennan et al., 2022).

The reduced speed of fitness recovery under multiple simultaneous stressors (Fig. 2) could reduce the time available for adaptation to reverse population decline before extinction occurs (Gomulkiewicz & Holt, 1995; Lindsey et al., 2013; Bell, 2017). If this slowdown of recovery applies broadly across species and stressor combinations, climate-based forecasts that extrapolate from single-stressor adaptive responses may overestimate the resilience of natural populations to realistic combinations of stressors.

Adding warming to the salinity decline selection regime also sharply reduced the extent of parallelism among the replicate lines, reducing the predictability of evolutionary outcomes (Fig. 5). As reported in the sections above, under salinity decline alone, each pair of replicate lines shared 23.8% of the selected haplotype blocks (mean pairwise Jaccard index) by Generation 20, whereas under the combined salinity and temperature selection regime each pair shared only 7.1% by Generation 25. Thus, under combined stressors, the reproducibility and consistency of the selection response will be lower. In addition, only 14% of candidate SNPs and 7.2% of enriched Gene Ontology terms were shared between the S and ST selection regimes, such that the well-characterized selection response to salinity decline alone (Stern et al., 2022; Du et al., 2026) provides a poor guide for predicting the response under salinity decline combined with warming. Thus, forecasts of adaptive capacity that rely on candidate loci from single-stressor studies, including genotype-environment association and genomic offset approaches (Waldvogel et al., 2019; Bernatchez et al., 2024), become less reliable under the multi-stressor conditions that natural populations are now experiencing.

More broadly, as anthropogenic climate change increasingly imposes simultaneous rather than single-axis environmental shifts, resolving how multiple simultaneous stressors reshape the genomic mechanisms of adaptation will be crucial for predicting which populations can achieve Evolutionary Rescue and which will likely fail to adapt in time.

## Methods

### Population sampling and experimental design

Wild individuals of the copepod *Eurytemora carolleeae* (Atlantic clade of the *E. affinis* species complex; Lee, 2000; Alekseev & Souissi, 2011) (haploid chromosome number = 4; Du et al., 2024) were collected in 2020 and 2021 from a saline marsh at Baie de L’Isle Verte in the St. Lawrence estuary, Quebec, Canada (48°00’16″N, 69°25’01″W), where salinity fluctuates between 5 and 30‰ (up to 40‰) (Winkler et al., 2008; Beyrend-Dur et al., 2009; Posavi et al., 2014; Stern & Lee, 2020; Rioux et al., 2023). Approximately 1,000 individuals were combined and maintained in the laboratory at 15‰ and at the native habitat temperature (12°C) for 3 to 4 generations to acclimate to laboratory conditions and to promote random mating and recombination. Each beaker initially contained ∼500 mixed-stage copepods derived from the acclimated population.

After laboratory acclimation, the population was subdivided into replicate lines. Seven selection lines were established for both the salinity decline (S) and salinity decline + temperature increase (ST) experiments (Fig. 1b), while six lines were established as control lines. One generation after subdivision into replicate lines, Generation 0 samples were collected from each selection and control line as the starting baseline by pooling 50 adults (25 males and 25 females) for sequencing (see below).

In the S experiment, selection lines were subjected to a stepwise salinity decline to impose natural selection for freshwater tolerance (Fig. 1b). Starting from 15‰, salinity was gradually reduced at each generation until reaching freshwater conditions (0‰, Lake Michigan water, ∼300 µS/cm conductivity) over six generations, following the salinity transition of 15 → 10 → 5 → 1 → 0.1 → 0.01 → 0‰. After reaching 0‰ at Generation 6, selection lines were maintained under freshwater conditions until Generation 20. In the ST experiment, selection lines experienced the same stepwise salinity decline simultaneously with temperature rise from 12°C to 22°C during the salinity transition phase (12 → 14 → 16 → 18 → 20 → 22 → 20°C). After the salinity and temperature transition, ST lines were maintained at 0‰ and 20°C until Generation 25 (Fig. 1b). Control lines were maintained at 15‰ and 12°C throughout the experiment (Fig. 1b).

Generations were not propagated by discrete transfers. Instead, individuals were maintained within the same beaker and allowed to reproduce continuously under ambient salinity and temperature conditions, resulting in overlapping generations. A generation time of approximately 3 weeks (21 days) was used to schedule salinity changes and sample collection (Katona, 1970; Lee et al., 2003). Because elevated temperatures accelerate copepod development, development time was modeled as a power function of temperature using nonlinear regression based on empirical data (Development time = 557.25 × Temperature^−1.272^; R² = 0.99) (Heinle & Flemer, 1975). Based on this relationship, a generation time of approximately 14 days was estimated for ST lines at our experimental temperatures above 18°C.

All lines were maintained on a diet of cryptophyte algae, with the marine alga *Rhodomonas salina* fed at moderate to high salinities (5–15‰) and the freshwater alga *R. minuta* at low salinities (≤ 1‰), to ensure adequate nutrition without osmotic shock to the algae (Vanderploeg et al., 1996; Tremblay et al., 2007). Culture water at 5–15‰ was prepared with Instant Ocean artificial sea salt and filtered deionized water with 20 mg/L Primaxin added to prevent bacterial infection. Water was changed weekly. Due to extinctions, 4 out of 7 S selection lines survived to Generation 20, and 6 out of 7 ST selection lines survived to Generation 25.

### Phenotypic measurements

Reproductive output was quantified as the number of eggs per clutch for gravid females at each sampling time point. Sampling generations were chosen to capture both the acute fitness decline during the salinity and temperature transition and the subsequent recovery phase (Generations 4, 5, 6, 8, 10 for S lines; Generations 4, 6, 11, 17 for ST lines; Generations 4, 6, 8, 10 for control lines). Sampling intervals differed between experiments because ST lines at elevated temperature had shorter generation times (estimated ∼14 days above 18°C versus ∼21 days at 12°C; see above), requiring adjusted sampling schedules to capture comparable developmental stages. At each generation sampled, 5 to 10 gravid females from each surviving replicate line were isolated, and their external egg sacs were removed. Eggs in the egg sacs were counted under a stereo microscope. Mean egg number (± SE) was calculated across all replicate lines within each experiment at each generation.

### DNA extraction, library preparation, and sequencing

Genomic DNA was extracted from pooled samples of 50 adults (25 males and 25 females) per line at multiple time points using a standard CTAB-based DNA extraction protocol (Du et al., 2024). For the S experiment and control lines, pooled samples were collected at Generations 0, 2, 3, 4, 5, 6, 7, 10, and 20. For the ST experiment lines, samples were collected at Generations 0, 2, 3, 4, 5, 6, 7, 15, and 25. Sequencing libraries were prepared with an average insert size of ∼350 bp using Nextera DNA Library Prep Kits (Illumina, San Diego, CA). Prepared samples were sequenced on an Illumina NovaSeq 6000 platform (2 × 150 bp reads) targeting ∼30× sequencing coverage per sample.

In total, the time-series dataset comprised pooled whole-genome sequences from 176 samples across both experiments and all time points (S experiment: 7 lines × 9 time points, minus three extinct samples = 60; ST experiment: 7 lines × 9 time points, minus one extinct sample = 62; Controls: 6 lines × 9 time points = 54).

### Sequence data processing and variant calling

Adapters and low-quality bases were trimmed using Fastp v0.23.0 (Chen et al., 2018). Clean reads were mapped to the chromosome-level reference genome of *E. carolleeae* (N = 4 chromosomes, 529 Mb) (Du et al., 2024) using BWA-MEM v0.7.12 (Li, 2013) with default parameters. Only confidently mapped and properly paired reads were retained, with duplicates removed using Picard v2.21.6. BAM files were sorted, converted to pileup format with SAMtools v1.21 (Li et al., 2009), discarding low-quality alignments and bases (Q < 20). SNPs were called using VarScan v2.4.6 (Koboldt et al., 2012) and filtered with BCFtools v1.21 (Danecek et al., 2021). The resulting VCF files were processed using the R package *poolfstat* v1.1.1 (Hivert et al., 2018), retaining only high-quality biallelic SNPs with minor allele frequency > 0.01, ≥ 4 reads per base call, and a total read depth of 10 to 200 across all pools. In total, 175,933 SNPs were identified, with 168,532 mapped onto the four assembled chromosomes of the *E. carolleeae* genome. For each SNP, allele frequencies were estimated at each time point for each line from the pooled sequencing read counts.

### Detection of signatures of selection

Multiple statistical tests were applied to identify SNPs showing allele frequency changes consistent with selection. Cochran–Mantel–Haenszel (CMH) tests were performed using the R package developed by Spitzer et al. (2020) to detect SNPs with frequency shifts exceeding expectations under random genetic drift across replicate lines. Chi-square tests were applied within each individual selection line using the same package. Both CMH and Chi-square test statistics were calibrated using estimated effective population sizes (*N*_e_) for each line (Table S14), estimated with poolSeq version 0.3.5 (Taus et al., 2017; Jonas et al., 2016). *P*-values were converted to *q*-values using the R package *qvalue* (Storey, 2003) to correct for multiple testing, and SNPs with *q* < 0.05 were considered significant.

To detect SNPs with frequency trajectories that diverged significantly between selection and control lines, we employed piecewise linear mixed models (LMMs) using the R package *lme4* (Bates et al., 2015). A single linear slope across all generations would not accurately capture the biphasic dynamics of allele frequency change, which showed minimal response during the salinity transition followed by a stronger response during the maintenance phase. In addition, the S and ST experiments had different sampling generations at the later phase of selection (i.e., Generations 10 and 20 in S lines versus Generations 15 and 25 in ST lines).

Therefore, allele frequency (*y*) was modeled using a piecewise linear specification with a breakpoint at generation *k* = 7, corresponding to the end of the salinity and temperature transition phase: *t_early* = min(Generation, *k*); *t_late* = max(Generation – *k*, 0). This piecewise specification accommodated the non-linear dynamics observed in both experiments and allowed the treatment effect to differ between the early (salinity and temperature transition) and late (maintenance and continuous selection) phases.

Two nested linear models were compared via likelihood ratio tests (LRT), fitted independently at each SNP. The Null model was: *y* ∼ *Treatment* + *t*_early + *t*_late + *Coverage* + (1|*Beaker*), whereas the Alternative model was: *y* ∼ *Treatment* × (*t*_early + *t*_late) + *Coverage* + (1|*Beaker*). The response variable *y* was the arcsine-square-root-transformed frequency of the focal allele at SNP *i* in replicate line *j* at generation *t*, where the focal allele was the one that rose in frequency in the selection lines (see above). Each observation therefore corresponded to one replicate line at one sampled generation. The fixed effects were *Treatment*, *t_early*, *t_late*, and *Coverage*, whereas *Beaker* was modeled as a random factor. *Treatment* was a two-level factor contrasting selection lines with control lines. The S and ST lines were each compared to the control lines in separate analyses. The factors *t_early* and *t_late* were the two continuous piecewise time periods defined above (previous paragraph), expressed in generations and partitioning each trajectory at the breakpoint *k* = 7. *Coverage* was included as a continuous covariate indicating the pooled sequencing read depth at that SNP in that sample. *Beaker* referred to the identity of the specific replicate of the selection or control lines and was fitted as a random intercept to account for line-to-line variation in baseline allele frequency.

Observations were weighted by the effective number of alleles sampled, calculated as *N*eff = (100 × *Coverage* – 1) / (100 + *Coverage*) (Feder et al., 2012), where 100 is the number of haploid genome copies per pool (50 diploid adults), and *Coverage* is the read depth at that SNP. The terms of interest were the *Treatment* × *t*_early and *Treatment* × *t*_late interactions, which tested whether the rate of allele frequency change differed between selection and control lines during the early and late phases, respectively. LRT statistics were compared to a Chi-square distribution with two degrees of freedom to obtain *P*-values, which were corrected to *q*-values (Storey, 2003).

### Identification of selected haplotype blocks

The union of candidate SNPs identified by the CMH, Chi-square, and LMM tests was used as input to the R package *haplovalidate* (Otte & Schlötterer, 2021b; Franssen et al., 2017b), which groups proximate candidate SNPs with correlated allele frequency shifts into putatively independent haplotype blocks. Each haplotype block was treated as a single “selected allele.” The median allele frequency of all SNPs within a haplotype block was used to represent the allele frequency of the haplotype block at each time point in each replicate line. A haplotype block was classified as under selection in a given replicate line if its frequency increase exceeded the 99.9th percentile of 10,000 neutral drift simulations performed using *poolSeq* (Taus et al., 2017), given the starting frequency of the haplotype block and the estimated *N*_e_. Allele frequencies were polarized so that the rising allele in the selection lines was tracked.

### Selection coefficient estimation

For each SNP or selected haplotype block, the selection coefficient (*s*) was estimated by fitting a linear mixed-effects model of the logit-transformed SNP frequency or the logit-transformed median allele frequency against generation using the R package *lme4* (Bates et al., 2015). The replicate line was treated as a random effect, and the observations were weighted by sequencing depth and *N*eff (effective number of alleles). The slope of this linear mixed-effects model provides an estimate of *s* under the assumption of approximately constant selection (Taus et al., 2017).

### Quantification of parallelism among replicate selection lines

Parallelism among replicate selection lines was quantified using the pairwise Jaccard index (Jaccard, 1912), computed as the number of shared selected haplotype blocks (classified as “selected” in both lines of a pair) divided by the total number of unique selected blocks in either line. The mean Jaccard index was computed across all pairwise comparisons among surviving replicate lines at each sampled generation.

In addition, principal component analysis (PCA) was conducted to assess the direction and magnitude of genomic responses among replicate lines. VCF files containing SNPs underlying selected haplotype blocks were generated. PCA was performed using PLINK v1.9 (Purcell et al., 2007) based on the allele frequency matrix, with each replicate line at each generation treated as a separate observation. The resulting principal components were further analyzed and visualized in R.

### Overlap of selected targets and functional enrichment in selection lines

Overlap in selected targets between the S and ST experiments was quantified at four levels of genomic organization. At the candidate SNP level, the number and proportion of SNPs identified by the combined CMH, Chi-square, and LMM tests that were shared between experiments were computed. At the level of SNPs contained within haplotype blocks, the same metrics were computed. At the genomic region (selected haplotype block) level, the Jaccard index of the base-pair coverage of selected haplotype blocks on each chromosome and genome-wide was calculated. At the gene level, genes underlying selected haplotype blocks were defined as those containing at least one selected-block SNP. The number, proportion, and Jaccard index of genes shared between the two experiments were computed. In addition, the genome-wide Spearman rank correlation was computed between the per-SNP selection coefficients of the two experiments across all 168,532 SNPs. Gene Ontology enrichment analyses were performed using TBtools v. 1.112 (Chen et al., 2020) for genes containing SNPs within selected haplotype blocks in each experiment.

## Acknowledgements

This work was funded by National Science Foundation grants IOS-2412790, DEB-2055356, OCE-1658517, and French National Research Agency ANR-19-MPGA-0004 (Macron’s “Make Our Planet Great Again” award) to Carol E. Lee. We thank members of the Lee laboratory for assistance with copepod culture maintenance and Grégoire Cortial and Gesche Winkler for collecting *Eurytemora carolleeae* copepod samples from Baie de L’Isle Verte (St. Lawrence estuary), Quebec, Canada.

## Author contributions

Conceptualization: C.E.L. and Z.D.; Methodology: C.E.L. and Z.D.; Formal analysis: Z.D.; Investigation: Z.D.; Resources: Z.D., A.T., L.L., S.L., and C.E.L.; Writing – original draft: Z.D. and C.E.L.; Writing – review & editing: Z.D., A.T., L.L., S.L., and C.E.L.; Visualization: Z.D. and C.E.L.; Supervision: C.E.L. and Z.D.; Project administration: C.E.L.; Funding acquisition: C.E.L.

## Data availability

The Pool-seq data generated in this study have been deposited in the NCBI Sequence Read Archive (SRA) under BioProject PRJNA1088474 (reviewer link: https://dataview.ncbi.nlm.nih.gov/object/PRJNA1088474?reviewer=ip7fc26shl1veidmjep9edtvv8). The reference genome of *E. carolleeae* used in this study is available in NCBI under BioProject PRJNA1075304 (https://www.ncbi.nlm.nih.gov/bioproject/PRJNA1075304) and at figshare (https://doi.org/10.6084/m9.figshare.29104271). Scripts for sequence data processing and selection analyses are available on Zenodo (https://doi.org/10.5281/zenodo.6615047). Allele frequency data (SNP and haplotype block) used in the statistical analysis are available at figshare.

