## Supplementary Figure for "Climate warming reduces the speed and predictability of polygenic adaptation to salinity decline"

**Table of Contents**

**Supplementary Figure 1.** Manhattan plot of SNPs under selection in the salinity (S) experiment lines on Chromosome 1 of the *Eurytemora carolleeae* genome.

**Supplementary Figure 2.** Manhattan plot of SNPs under selection in the salinity (S) experiment lines on Chromosome 2 of the *Eurytemora carolleeae* genome.

**Supplementary Figure 3.** Manhattan plot of SNPs under selection in the salinity (S) experiment lines on Chromosome 3 of the *Eurytemora carolleeae* genome.

**Supplementary Figure 4.** Manhattan plot of SNPs under selection in the salinity (S) experiment lines on Chromosome 4 of the *Eurytemora carolleeae* genome.

**Supplementary Figure 5.** Manhattan plot of SNPs under selection in the salinity + temperature (ST) experiment lines on Chromosome 1 of the *Eurytemora carolleeae* genome.

**Supplementary Figure 6.** Manhattan plot of SNPs under selection in the salinity + temperature (ST) experiment lines on Chromosome 2 of the *Eurytemora carolleeae* genome.

**Supplementary Figure 7.** Manhattan plot of SNPs under selection in the salinity + temperature (ST) experiment lines on Chromosome 3 of the *Eurytemora carolleeae* genome.

**Supplementary Figure 8.** Manhattan plot of SNPs under selection in the salinity + temperature (ST) experiment lines on Chromosome 4 of the *Eurytemora carolleeae* genome.

**Supplementary Figure 9.** Limited and uneven overlap of genomic targets between the salinity (S) and salinity + temperature (ST) experiments.

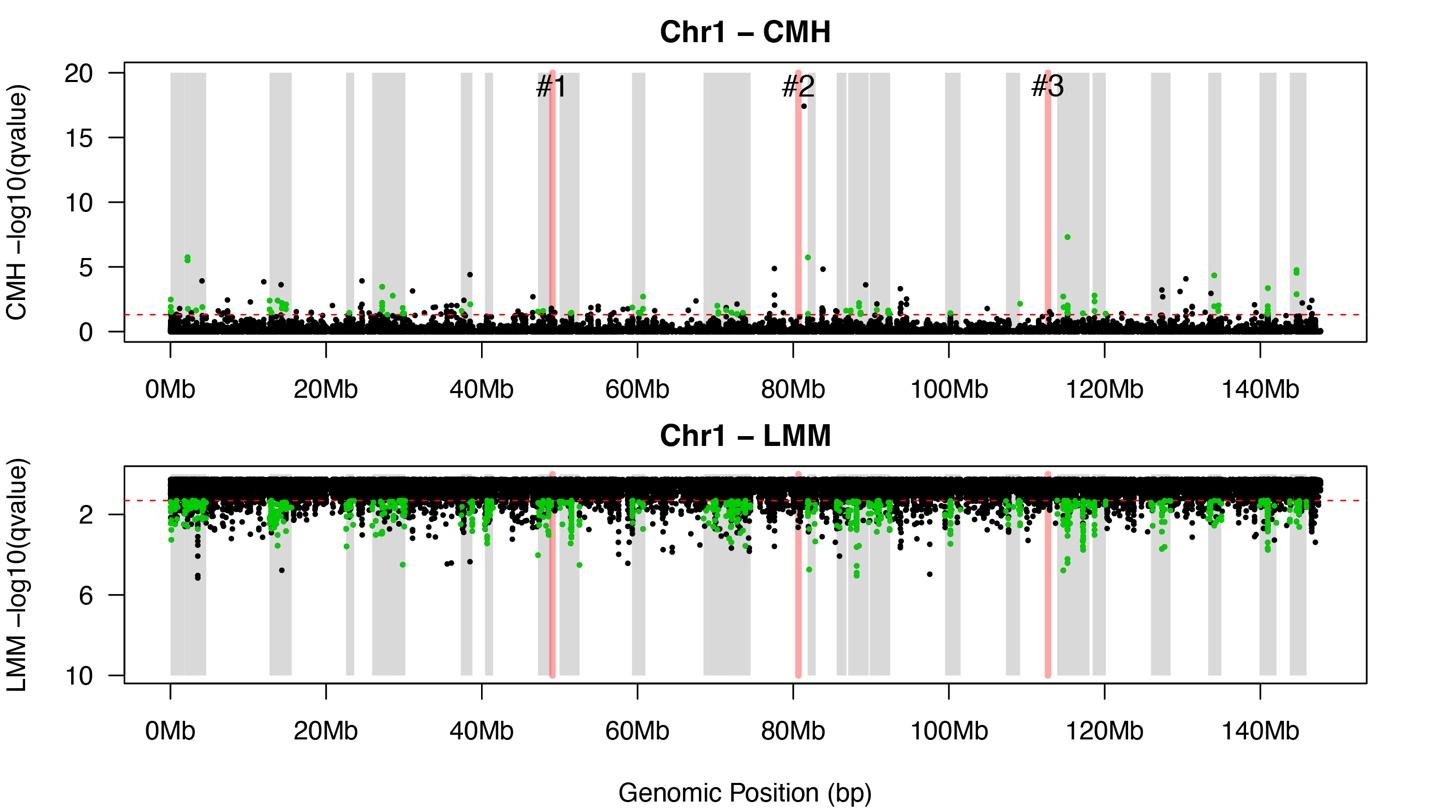

**Supplementary Figure 1.** **Manhattan plot of SNPs under selection in the salinity (S) experiment lines on Chromosome 1 of the *Eurytemora carolleeae* genome.** SNPs beyond the dotted red line were deemed significant after correction for multiple testing (*q* < 0.05). Shaded gray bars delineate haplotype blocks identified as targets of selection on this chromosome. Shaded red bars delineate positions of chromosomal fusion sites. Top: results of the Cochran–Mantel–Haenszel (CMH) test, detecting significant allele frequency changes beyond expectations from genetic drift. Bottom: results of the linear mixed model (LMM) test, distinguishing allele frequency trajectories between selection and control lines. Both the top and bottom panels show candidate SNPs under selection in the selection lines, detected by two different tests.

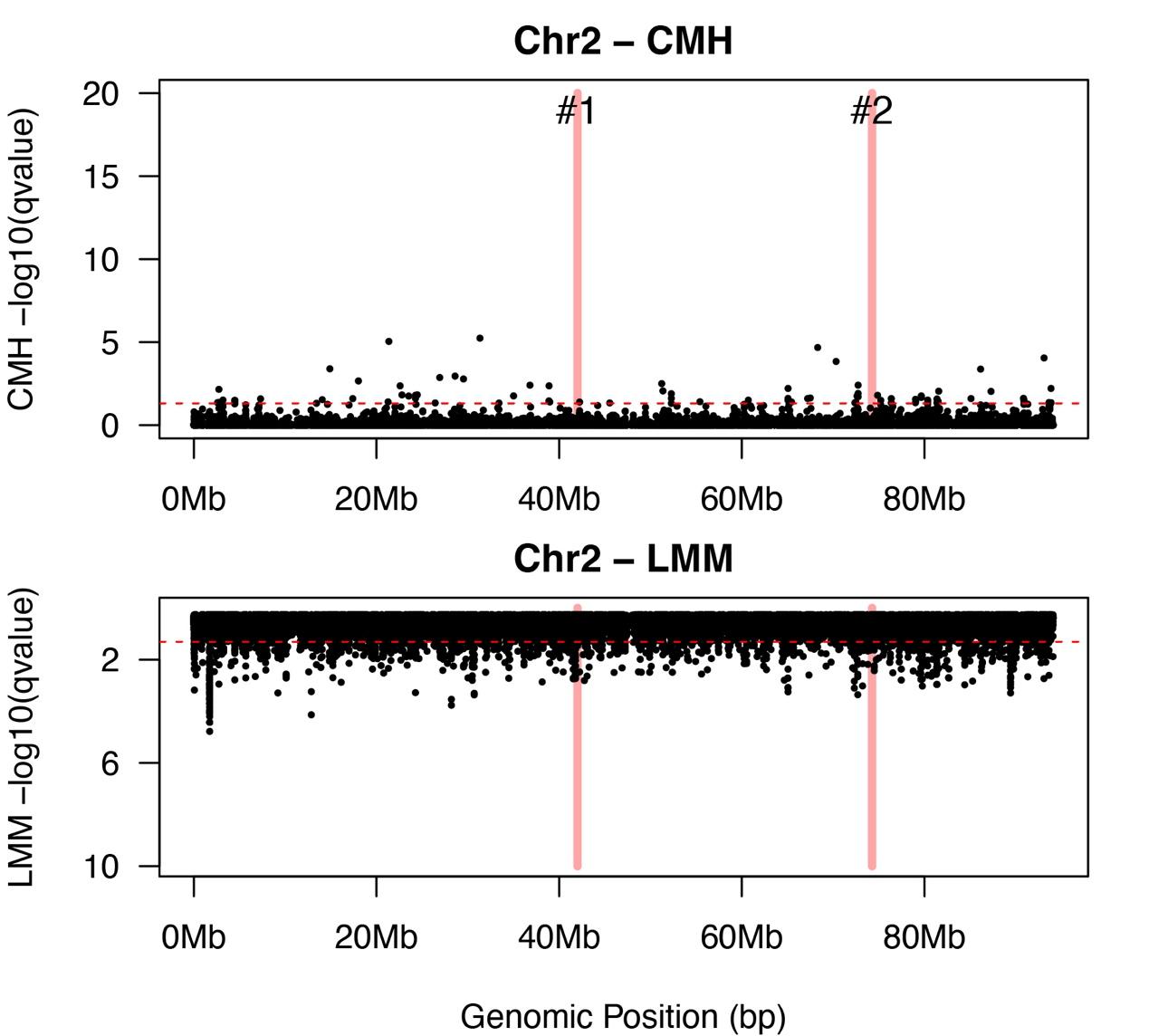

**Supplementary Figure 2.** **Manhattan plot of SNPs under selection in the salinity (S) experiment lines on Chromosome 2 of the *Eurytemora carolleeae* genome.** SNPs beyond the dotted red line were deemed significant after correction for multiple testing (*q* < 0.05). Shaded gray bars delineate haplotype blocks identified as targets of selection on this chromosome. Shaded red bars delineate positions of chromosomal fusion sites. Top: results of the Cochran–Mantel–Haenszel (CMH) test, detecting significant allele frequency changes beyond expectations from genetic drift. Bottom: results of the linear mixed model (LMM) test, distinguishing allele frequency trajectories between selection and control lines. Both the top and bottom panels show candidate SNPs under selection in the selection lines, detected by two different tests.

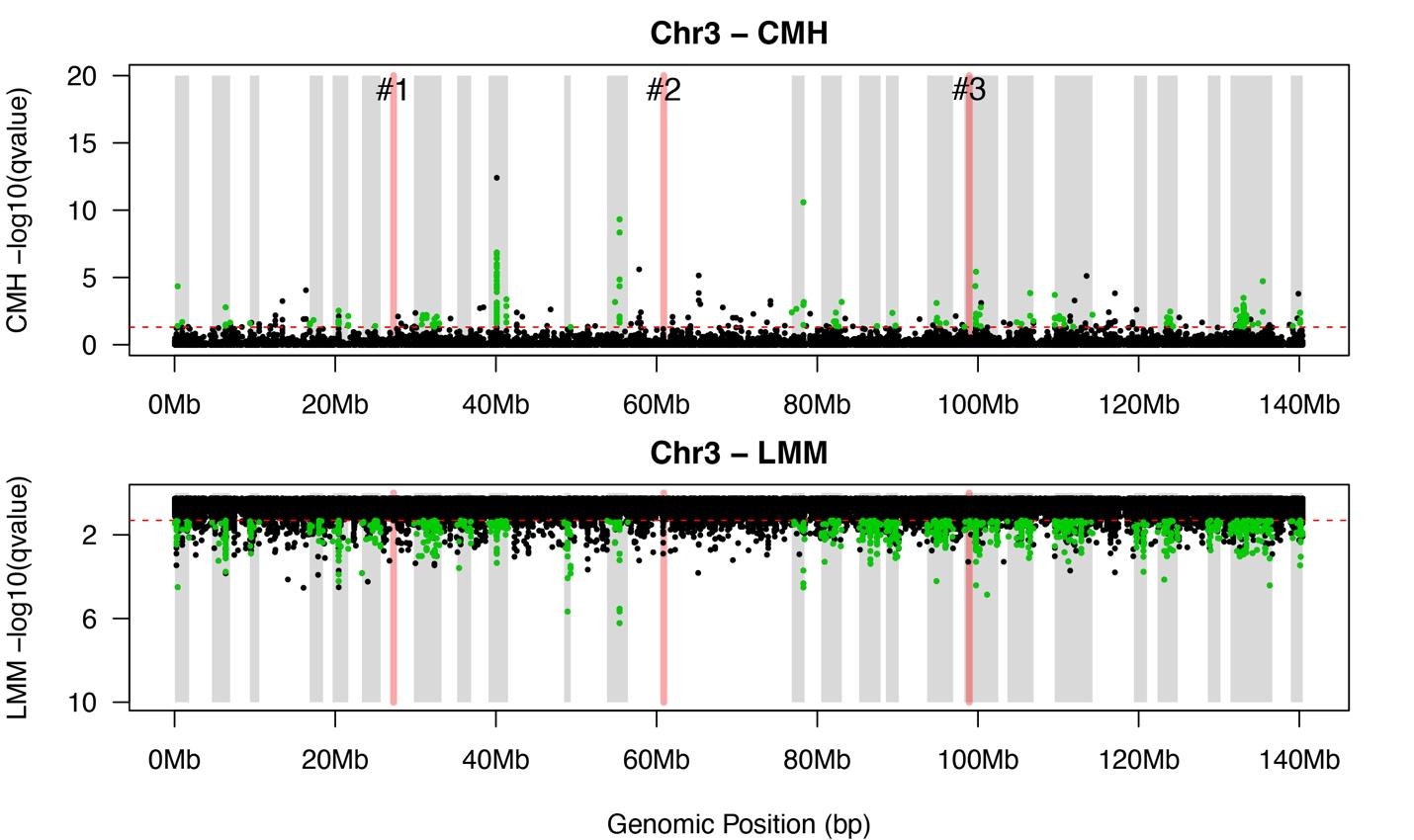

**Supplementary Figure 3.** **Manhattan plot of SNPs under selection in the salinity (S) experiment lines on Chromosome 3 of the *Eurytemora carolleeae* genome.** SNPs beyond the dotted red line were deemed significant after correction for multiple testing (*q* < 0.05). Shaded gray bars delineate haplotype blocks identified as targets of selection on this chromosome. Shaded red bars delineate positions of chromosomal fusion sites. Top: results of the Cochran–Mantel–Haenszel (CMH) test, detecting significant allele frequency changes beyond expectations from genetic drift. Bottom: results of the linear mixed model (LMM) test, distinguishing allele frequency trajectories between selection and control lines. Both the top and bottom panels show candidate SNPs under selection in the selection lines, detected by two different tests.

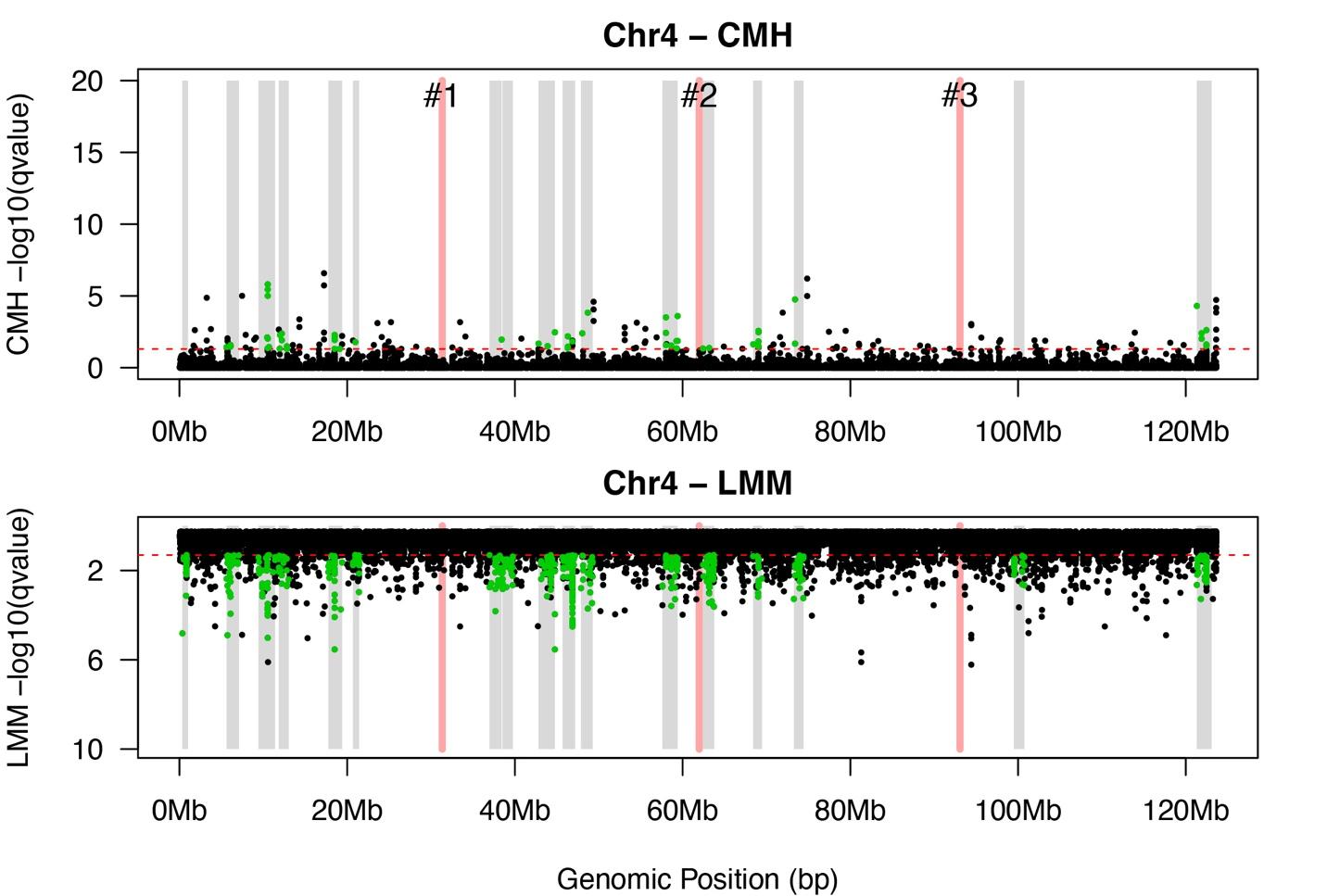

**Supplementary Figure 4.** **Manhattan plot of SNPs under selection in the salinity (S) experiment lines on Chromosome 4 of the *Eurytemora carolleeae* genome.** SNPs beyond the dotted red line were deemed significant after correction for multiple testing (*q* < 0.05). Shaded gray bars delineate haplotype blocks identified as targets of selection on this chromosome. Shaded red bars delineate positions of chromosomal fusion sites. Top: results of the Cochran–Mantel–Haenszel (CMH) test, detecting significant allele frequency changes beyond expectations from genetic drift. Bottom: results of the linear mixed model (LMM) test, distinguishing allele frequency trajectories between selection and control lines. Both the top and bottom panels show candidate SNPs under selection in the selection lines, detected by two different tests.

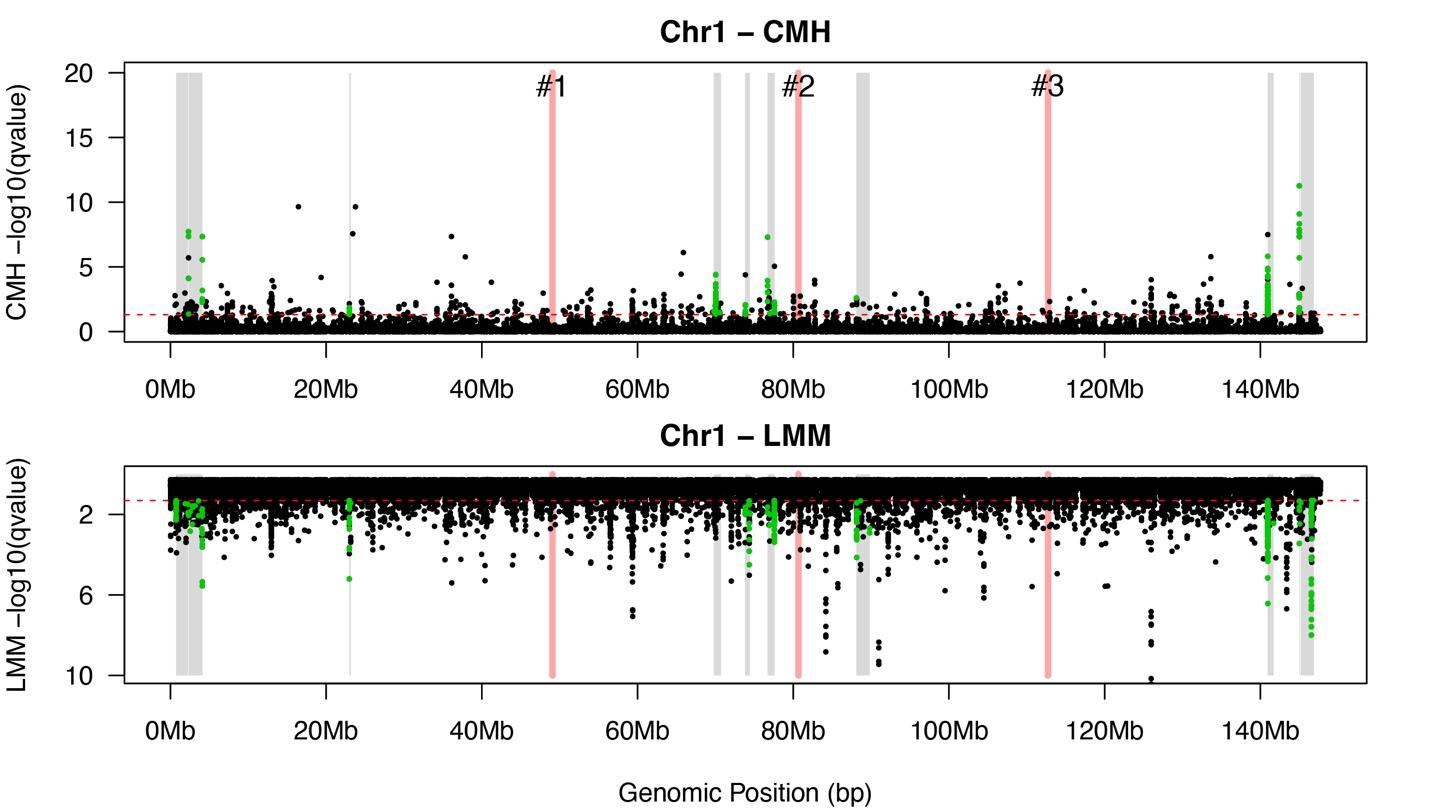

**Supplementary Figure 5.** **Manhattan plot of SNPs under selection in the salinity + temperature (ST) experiment lines on Chromosome 1 of the *Eurytemora carolleeae* genome.** SNPs beyond the dotted red line were deemed significant after correction for multiple testing (*q* < 0.05). Shaded gray bars delineate haplotype blocks identified as targets of selection on this chromosome. Shaded red bars delineate positions of chromosomal fusion sites. Top: results of the Cochran–Mantel–Haenszel (CMH) test, detecting significant allele frequency changes beyond expectations from genetic drift. Bottom: results of the linear mixed model (LMM) test, distinguishing allele frequency trajectories between selection and control lines. Both the top and bottom panels show candidate SNPs under selection in the selection lines, detected by two different tests.

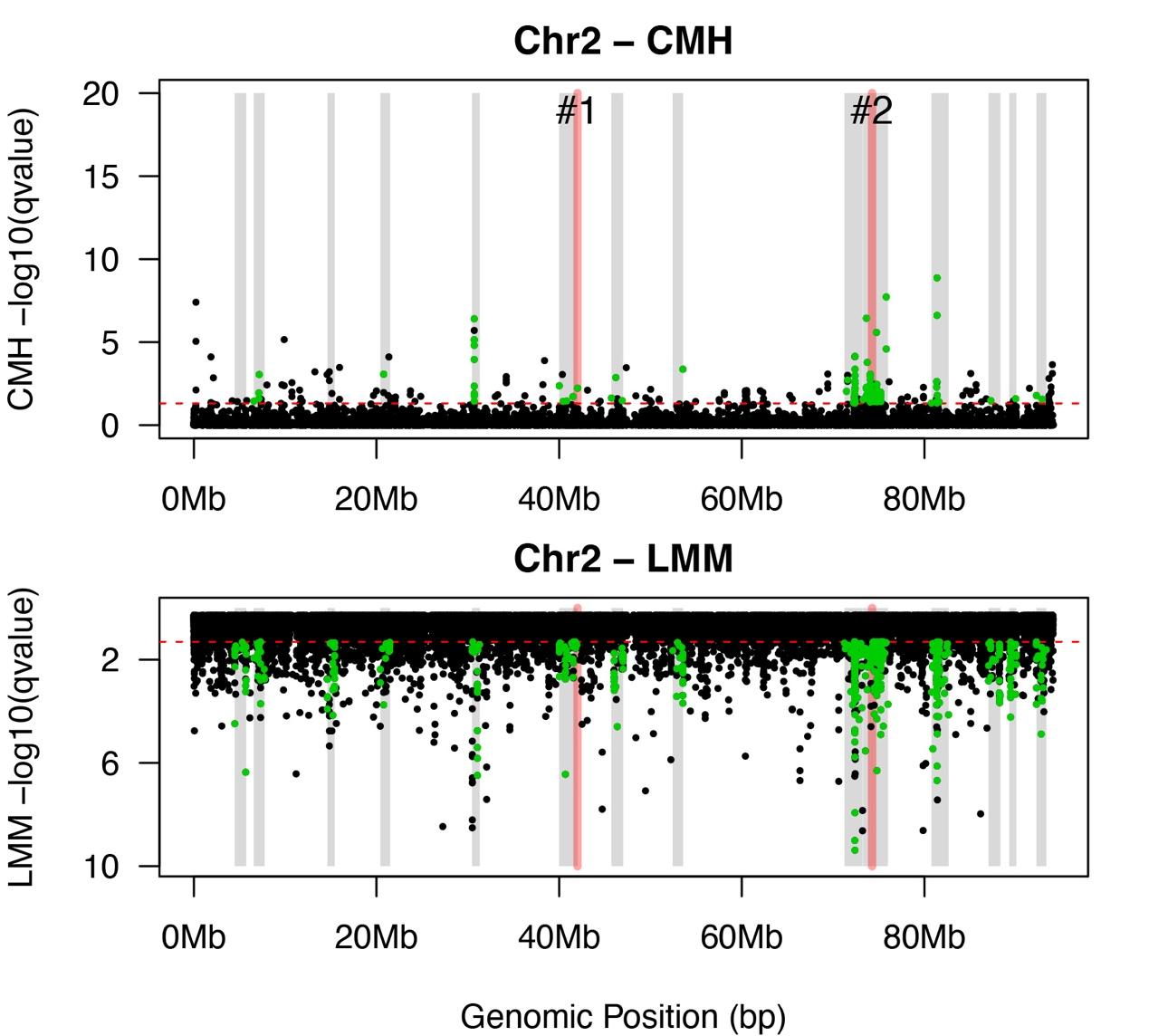

**Supplementary Figure 6.** **Manhattan plot of SNPs under selection in the salinity + temperature (ST) experiment lines on Chromosome 2 of the *Eurytemora carolleeae* genome.** SNPs beyond the dotted red line were deemed significant after correction for multiple testing (*q* < 0.05). Shaded gray bars delineate haplotype blocks identified as targets of selection on this chromosome. Shaded red bars delineate positions of chromosomal fusion sites. Top: results of the Cochran–Mantel–Haenszel (CMH) test, detecting significant allele frequency changes beyond expectations from genetic drift. Bottom: results of the linear mixed model (LMM) test, distinguishing allele frequency trajectories between selection and control lines. Both the top and bottom panels show candidate SNPs under selection in the selection lines, detected by two different tests.

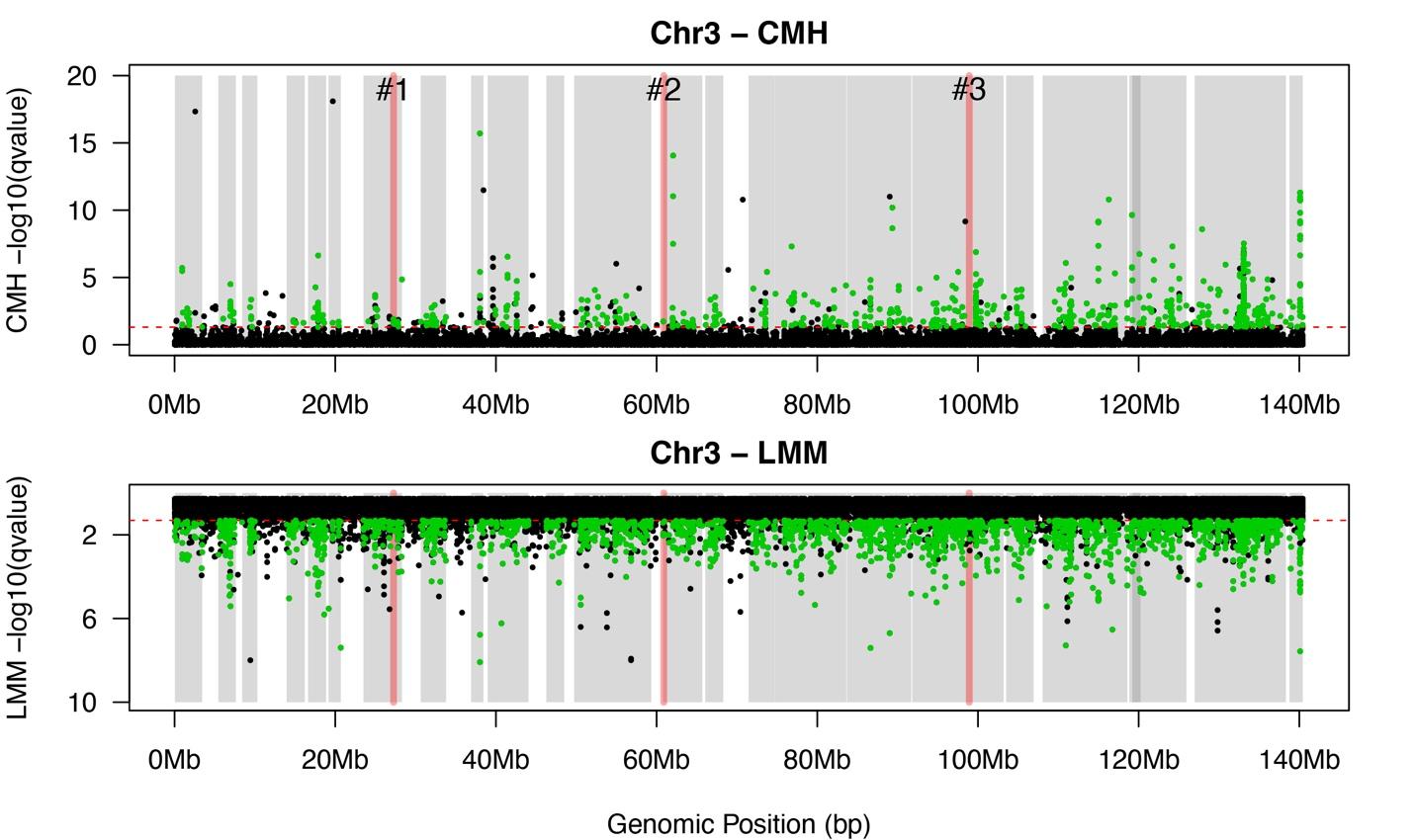

**Supplementary Figure 7.** **Manhattan plot of SNPs under selection in the salinity + temperature (ST) experiment lines on Chromosome 3 of the *Eurytemora carolleeae* genome.** SNPs beyond the dotted red line were deemed significant after correction for multiple testing (*q* < 0.05). Shaded gray bars delineate haplotype blocks identified as targets of selection on this chromosome. Shaded red bars delineate positions of chromosomal fusion sites. Top: results of the Cochran–Mantel–Haenszel (CMH) test, detecting significant allele frequency changes beyond expectations from genetic drift. Bottom: results of the linear mixed model (LMM) test, distinguishing allele frequency trajectories between selection and control lines. Both the top and bottom panels show candidate SNPs under selection in the selection lines, detected by two different tests.

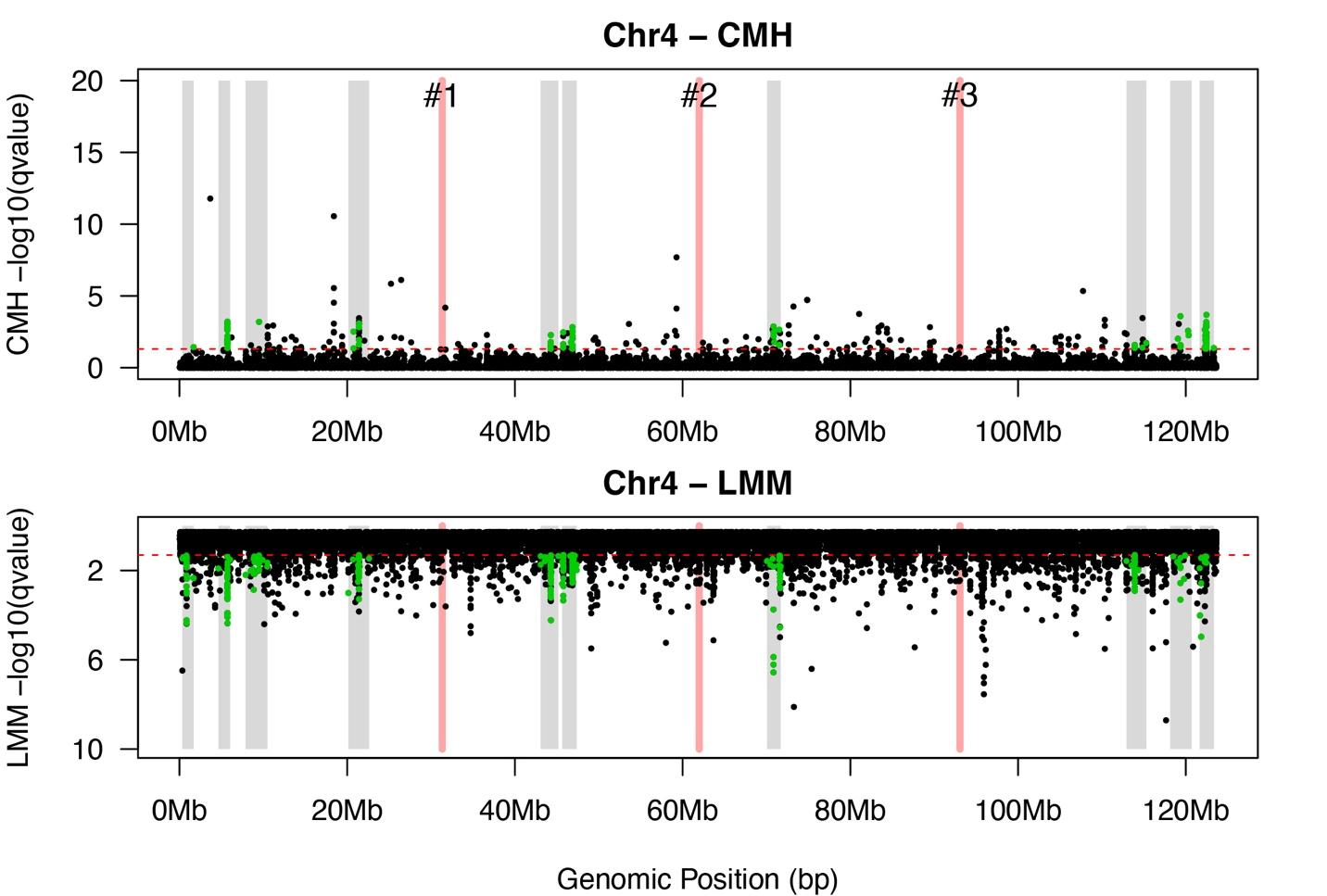

**Supplementary Figure 8.** **Manhattan plot of SNPs under selection in the salinity + temperature (ST) experiment lines on Chromosome 4 of the *Eurytemora carolleeae* genome.** SNPs beyond the dotted red line were deemed significant after correction for multiple testing (*q* < 0.05). Shaded gray bars delineate haplotype blocks identified as targets of selection on this chromosome. Shaded red bars delineate positions of chromosomal fusion sites. Top: results of the Cochran–Mantel–Haenszel (CMH) test, detecting significant allele frequency changes beyond expectations from genetic drift. Bottom: results of the linear mixed model (LMM) test, distinguishing allele frequency trajectories between selection and control lines. Both the top and bottom panels show candidate SNPs under selection in the selection lines, detected by two different tests.

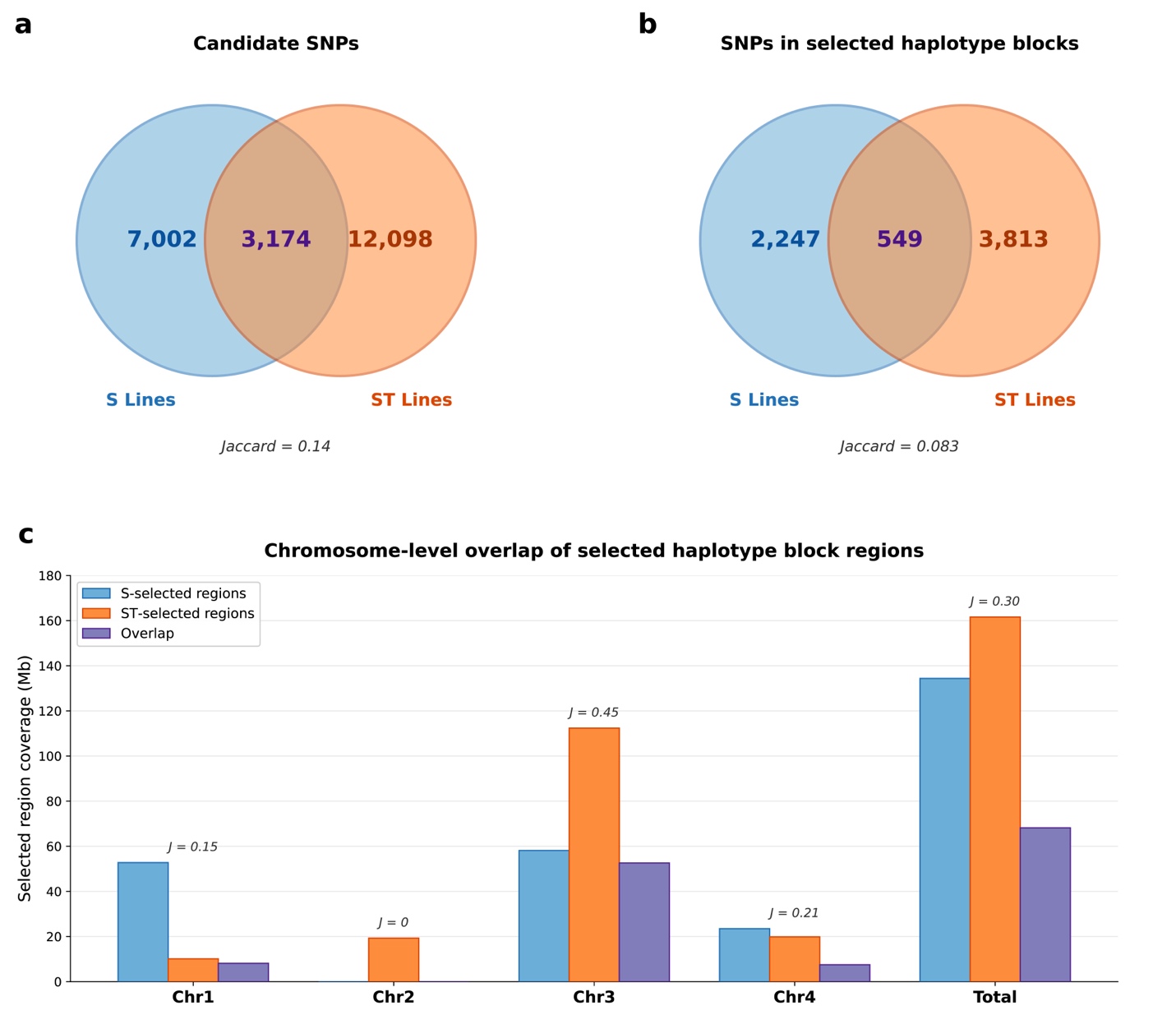

**Supplementary Figure 9. Limited and uneven overlap of genomic targets between the salinity (S) and salinity + temperature (ST) experiments.** (**a**) Venn diagram showing the overlap of candidate SNPs identified by the combined CMH, Chi-square, and LMM tests between the S (blue; 10,176 SNPs) and ST (orange; 15,272 SNPs) experiments. Only 3,174 candidate SNPs (Jaccard index = 0.14) were shared between experiments. (**b**) Venn diagram showing the overlap of SNPs within selected haplotype blocks between the S (blue; 2,796 SNPs on 66 blocks) and ST (orange; 4,362 SNPs on 58 blocks) experiments. Only 549 haplotype block SNPs (Jaccard index = 0.083) were shared, representing 19.6% of S and 12.6% of ST haplotype block SNPs. (**c**) Chromosome-level comparison of selected haplotype block region coverage (Mb) in the S experiment (blue), ST experiment (orange), and their overlap (purple). Jaccard indices (*J*) for each chromosome are shown above the bars. Chromosome 3 showed the highest level of concordance (*J* = 0.45). Chromosome 2 showed exclusively ST-specific selection (*J* = 0). Chromosomes 1 and 4 showed intermediate levels of overlap (*J* = 0.15 and 0.21, respectively). Genome-wide overlap was 0.30.
